# Dosage differences in *12-OXOPHYTODIENOATE REDUCTASE* genes modulate wheat primary root growth

**DOI:** 10.1101/2022.08.25.505338

**Authors:** G. Gabay, H. Wang, J. Zhang, J. I. Moriconi, G. F. Burguener, T. Howell, A. Lukaszewski, B. Staskawicz, M.-J. Cho, J. Tanaka, T. Fahima, H. Ke, K. Dehesh, G.-L. Zhang, J.-Y. Gou, M. Hamberg, G. Santa Maria, J. Dubcovsky

## Abstract

Wheat is an essential crop for global food security and is well adapted to a wide variety of soils^1^. However, the gene networks regulating different root architectures remain poorly understood. We report here the identification of a cluster of a monocot-specific *12-OXOPHYTODIENOATE REDUCTASE* genes from subfamily III (*OPRIII*) that modulate key differences in wheat root architecture associated with grain yield under water-limited conditions. Wheat plants with a loss-of-function mutation in *OPRIII* showed longer seminal roots, whereas plants with increased *OPRIII* dosage or transgenic over-expression showed reduced seminal root growth, precocious development of lateral roots and increased jasmonic acid (JA). A JA-biosynthesis inhibitor eliminated the root differences, confirming a JA-mediated mechanism. Multiple transcriptome analysis of transgenic and wild-type lines revealed significant enriched JA-biosynthetic and reactive oxygen species (ROS) pathways that paralleled changes in ROS distribution. The *OPRIII* genes provide a useful entry point to engineer root architecture in wheat and other cereals.

---

Further increases in wheat grain yield are required to feed a growing population, but losses generated by water stress are increasing with global warming, and are eroding progress in other areas of wheat improvement ^1^. Root depth and biomass distribution in the soil profile are critical traits for adaptation to water stress and have been prioritized for improving drought resilience in wheat ^2–4^. However, the genetic mechanisms that regulate root architecture in wheat remain largely unknown. This has prompted new efforts to understand and improve wheat root architecture to optimize water acquisition in both common (*Triticum aestivum*, genomes AABBDD) and durum wheat (*T. turgidum* ssp. *durum*, genomes AABB).

A tested source for improving these traits in wheat is the introgression of the short arm of rye (*Secale cereale* L.) chromosome one (1RS) into common wheat (henceforth 1RS.1BL), which induces higher root biomass and confers a yield advantage under drought stress ^5–8^. Unfortunately, this translocation reduces bread baking quality ^9,10^. To eliminate the negative effect on quality, a recombinant 1RS chromosome with two 1BS wheat interstitial introgressions (henceforth 1WW) was developed twenty years ago in the wheat cultivar Pavon ^11^. We introgressed the engineered chromosome into the 1RS.1BL cv. Hahn, and generated lines with the complete 1RS (1RS), the proximal (1WR) or the distal wheat introgression (1RW, Fig. 1A).

**Fig. 1.**
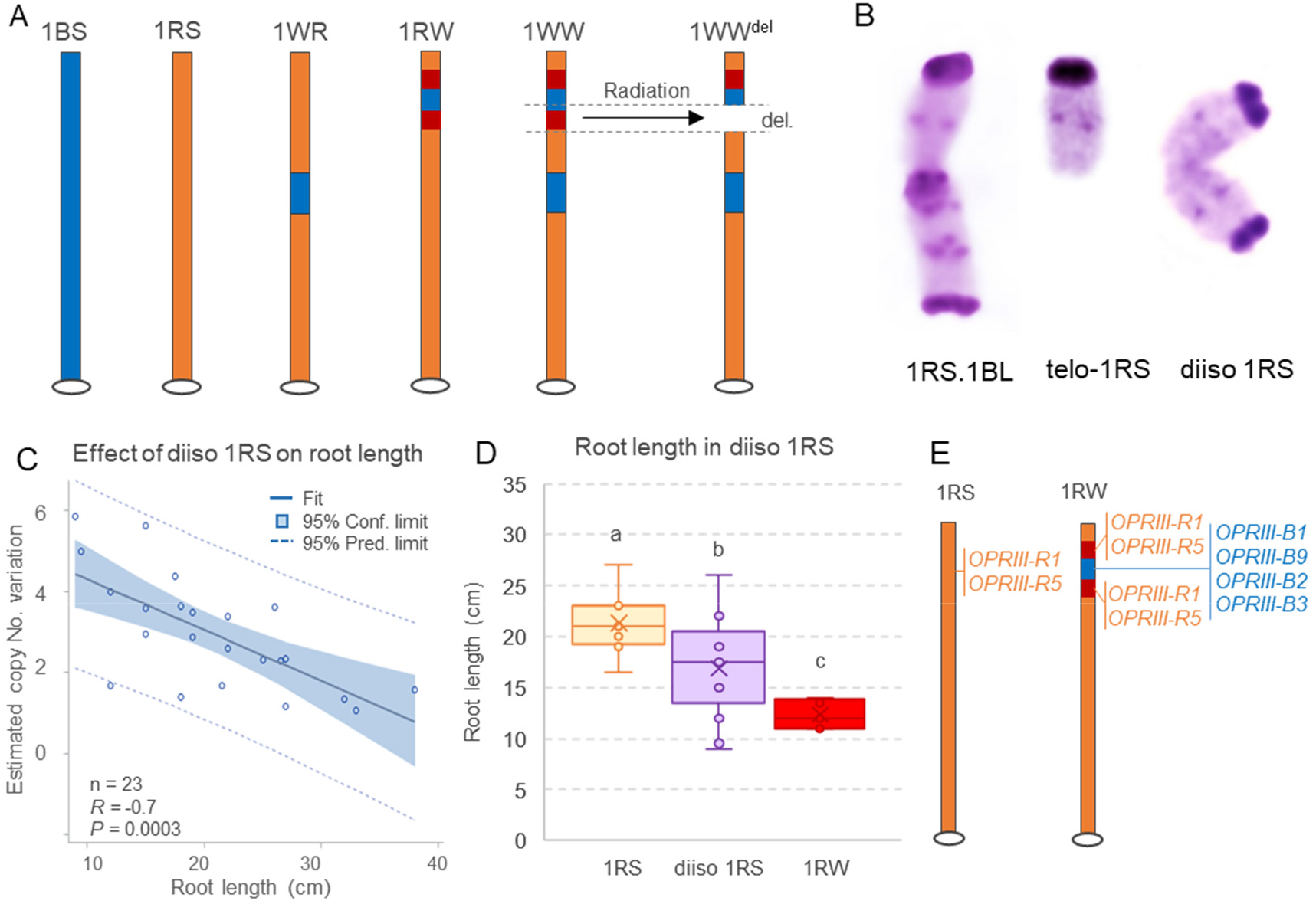
Wheat-rye genetic stocks and their effects on seminal root length. (**A**) Genetic stocks used in this project. The hexaploid wheat variety Hahn has a complete 1RS arm replacing the 1BS arm. Line 1WR has a proximal wheat segment introgressed in the 1RS short arm, whereas 1RW has a distal wheat segment introgression. Line 1WW has both wheat introgressions and 1WW^del^ has a 4.9 Mb deletion including part of the distal wheat introgression. Wheat chromatin is indicated in blue and rye in orange. The duplicated region in rye flanking the 1BS insertion is indicated in red. (**B**) C-banding of the 1RS.1BL translocation, the 1RS telocentric and a diiso 1RS chromosome. Hahn-1RS plants homozygous for diiso 1RS have four extra 1RS arms and, therefore, triple dosage of genes in the duplicated region. (**C**) Regression between 1RS copy number variation) and root length in hydroponic tanks 19 days after germination (n = 23). CNV of the 1RS arm was significantly correlated to the length of the seminal roots. (**D**) Seminal root length of 13 diiso 1RS lines with at least an extra 1RS arm (total 3-6 1RS arms, average 4.0 ± 0.3) relative 1RS plants (n= 8, two 1RS arms) and 1RW plants with a triplicated 1RS/1BS candidate gene region (n= 8, total 6 1RS copies). Different letters indicate significant differences in Tukey test (*P* < 0.05). Raw data and statistics are in Data S1. CNV, determined with primers qrt-1RS6.4 (Data S7) (**E**) Schematic representation of 1RS and 1RW showing duplicated *OPRIII* genes in 1RW.

In multi-year replicated field studies, lines carrying the complete 1RS arm showed roots reaching deeper layers of the soil, higher carbon isotope discrimination, and increased stomatal conductance, relative to 1RW. This resulted in improved canopy water status and up to 40% higher grain yields under terminal water stress ^12^. Under hydroponic conditions, elongation of the 1RW seminal roots decreased from nine days after germination (DAG) and almost stopped by ~16 DAG, and lateral roots formed close to the seminal root apical meristem (RAM). None of these phenotypes was observed in sister lines with the complete 1RS arm ^13^.

1RW chromosome has a complex structure generated by structural differences between the 1RS and 1BS distal chromosome regions ^14–16^. This chromosome has a duplicated 1RS region (7.0 Mb) flanking a colinear insertion from wheat chromosome arm 1BS (4.8 Mb), which resulted in increased dosage of the genes included in the triplicated region ^16^ (Fig. 1A). Other recombinant lines with non-duplicated 1BS or 1RS distal regions have long seminal roots, suggesting that changes in gene dosage were responsible for the shorter seminal roots. This hypothesis was supported by the intermediate seminal-root length of heterozygous 1RW plants ^16^. In this study we developed a diisosomic 1RS addition line with two fused 1RS arms into Hahn-1RS (diiso 1RS, Fig. 1B) to validate the dosage effect hypothesis. The seminal roots from plants carrying extra 1RS arms were significantly shorter than those from the wildtype (Fig. 1C-D, Data S1), confirming that increased gene dosage results in shorter seminal roots.

A radiation-induced deletion (Fig. 1A), resulted in the elimination of two of the three triplicated regions ^16^. This radiation mutant has long seminal roots similar to Hahn-1RS, demonstrating the importance of gene dosage, and delimiting the candidate gene region to the deleted segment. Among the 14 high-confidence genes annotated in the deleted 1BS region and expressed in the seminal roots ^16^, we prioritized four *OPRIII* genes. These genes encode enzymes involved in the biosynthesis of JA ^17^, whose JA-Ile conjugate is an active phytohormone proposed to modify root architecture and responses to drought in Arabidopsis ^18–20^ and rice ^21^.

The *OPR* gene family expanded in the vascular plants, where five well-conserved subfamilies (I-V) have been reported ^22,23^. The *OPR* genes described here belong to the monocot-specific subfamily III and have been designated as *12-OXOPHYTODIENOATE REDUCTASE SUBFAMILY III* (*OPRIII*) in a phylogenetic study of the *OPR* genes ^23^ present in the wheat genome ^24^. Gene names, coordinates and predicted proteins used in this study are described in Data S2. In rice, six tightly linked *OPR* genes from the same subfamily (*OPR6.1-OPR6.6*) have been described on the short arm of chromosome 6 ^17^, in a region colinear with wheat chromosome arm 7S that includes the linked *OPRIII12* and *OPRIII13* genes ^23^. These two wheat genes are more similar to the rice genes than to a cluster of 4-5 linked *OPRIII* genes located within the region of chromosome arm 1S that is triplicated in 1RW. Based on the phylogenetic tree in Extended Data Fig. S1 the rye genes were designated as *OPRIII-R2* and *OPRIII-R5*. The wheat region on chromosome arm 1S shows no colinear *OPR* genes in rice suggesting that the *OPRIII* expansion on 1S occurred in the Triticeae lineage. Wheat has additional *OPRIII* genes in other four chromosome regions ^23^.

A transcriptome analysis of the seminal root terminal region (1 cm) in 1RS and 1RW at 6 and 16 DAG (Data S3-5) revealed that most *OPRIII* genes are expressed at significantly higher levels at 6 DAG than at 16 DAG, a result confirmed by qRT-PCR (Extended Data Fig. S2). As expected, the duplicated rye *OPRIII* genes showed higher expression in 1RW than in 1RS (Extended Data Fig. S2), and *OPRIII-B1* (present in the 1BS introgression, Fig. 1E) was detected only in 1RW (Extended Data Fig. S2, Data S6-7). These changes contributed to a higher *OPRIII* expression in 1RW than in 1RS.

To validate the role of the *OPRIII* genes on seminal root length, we generated a 32-bp frame-shift CRISPR induced deletion in *OPRIII-B1(*mut-OPRIII-B1) in 1RW. Vectors used for the transgenic plants are described in Extended Data Fig. S3. The seminal roots of T_2_ plants without the transgene and homozygous for the deletion were 28% longer than the sister 1RW plants carrying the wildtype *OPRIII-B1* (*P*< 0.0001, Fig. 2A, Data S8). A replicated experiment using T_3_ plants showed similar results (Extended Data Fig. S4), demonstrating that reduced *OPRIII* dosage is associated with increased seminal root length in 1RW.

**Fig. 2.**
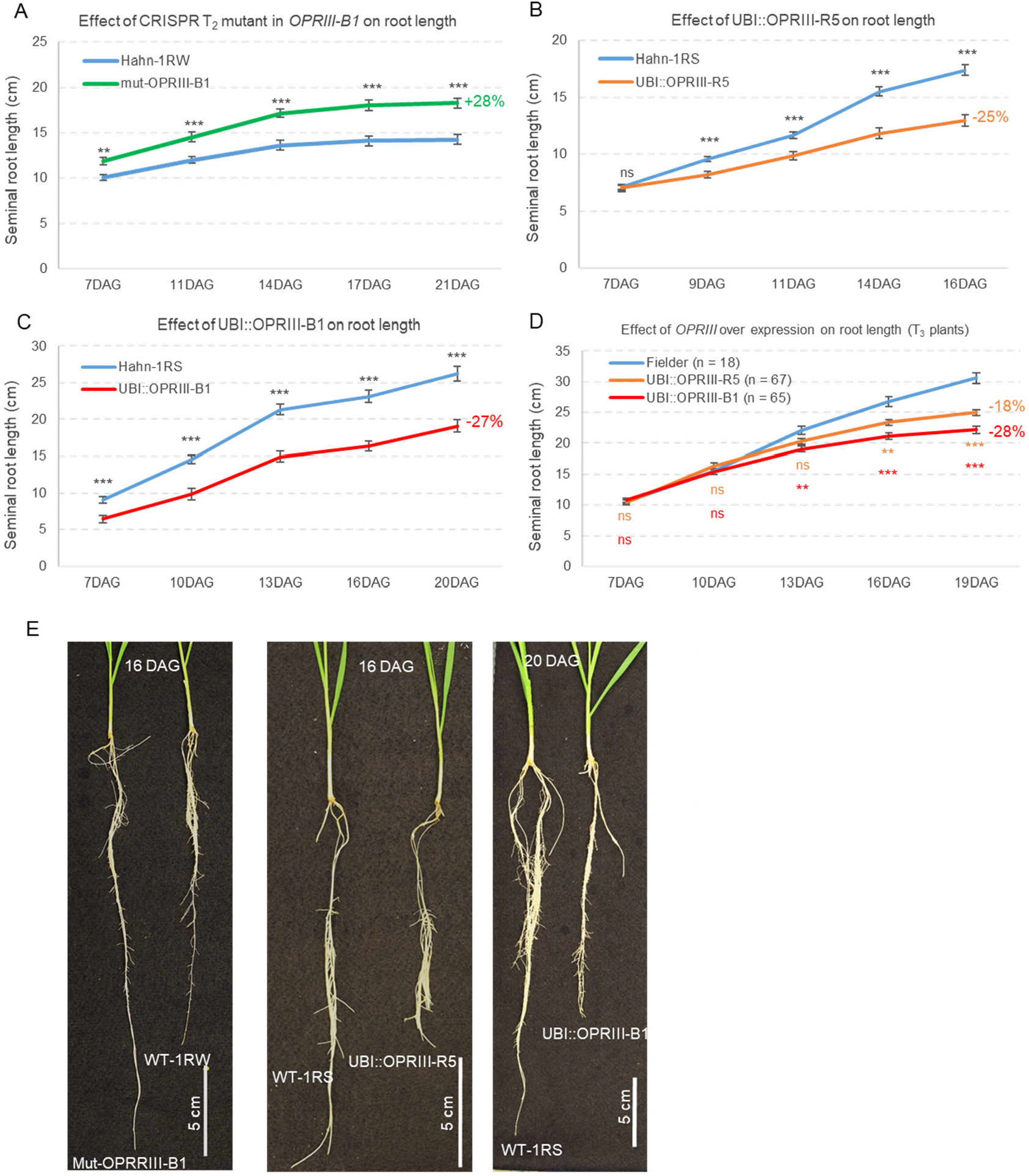
Effect of loss-of-function mutations and over-expression of *OPRIII* genes on root length. (**A**) Root length time course for 1RW T_2_ sister lines with and without a 32 bp deletion in *OPRIII-B1.* (**B-C**) Transgenic lines in a Hahn-1RS background constitutively expressing UBI::OPRIII-R5 (**B**) or UBI::OPRIII-B1 (**C**). (**D**) Transgenic lines in a Fielder background expressing the same two constructs **(E)** Images of roots lengths in the same genotype comparisons as in A to C. Raw data and statistical analyses are available in Data S8-11.

We then developed transgenic Hahn-1RS and Fielder cultivars constitutively expressing either wheat *OPRIII-B1* or rye *OPRIII-R5* genes driven by the maize *UBIQUITIN* promoter (Extended Data Fig. S3); and confirmed that both transgenes were expressed in leaves, where they were not detectable in the non-transgenic controls. At the end of the experiments, the seminal roots of the transgenic plants were 18-27% shorter than those in their non-transgenic sister lines (Fig. 2B-D and Extended Data Fig. S5). These differences varied in time resulting in highly significant genotype x time interactions in the repeated measure ANOVAs (Data S9-11), suggesting that the effects of constitutive expression of the *OPRIII* transgenes on seminal root elongation vary with root development.

In both UBI::OPRIII-R5 and 1RW, once seminal roots growth was arrested, secondary roots emerged significantly closer to the RAM than in 1RS (Fig. 3A-E, Data S12). This secondary roots showed an earlier growth arrest in UBI::OPRIII-R5 than in RW, with formation of lateral roots close to the secondary root’s RAM and a more extreme phenotype (Fig. 3A and E). Staining of these roots with nitro blue tetrazolium (NBT) revealed that the ROS were more restricted to the distal root region in 1RW and UBI::OPRIII-R5 than in 1RS (Fig. 3E), as previously described for 1RW ^13^. The significant effects of the loss-of-function mutation and over-expression of *OPRIII* genes on seminal root length and architecture demonstrate that the *OPRIII* genes are responsible for the different root phenotypes in 1RS and 1RW isogenic lines, and likely for their different agronomic performance under water stress ^12^.

**Fig. 3.**
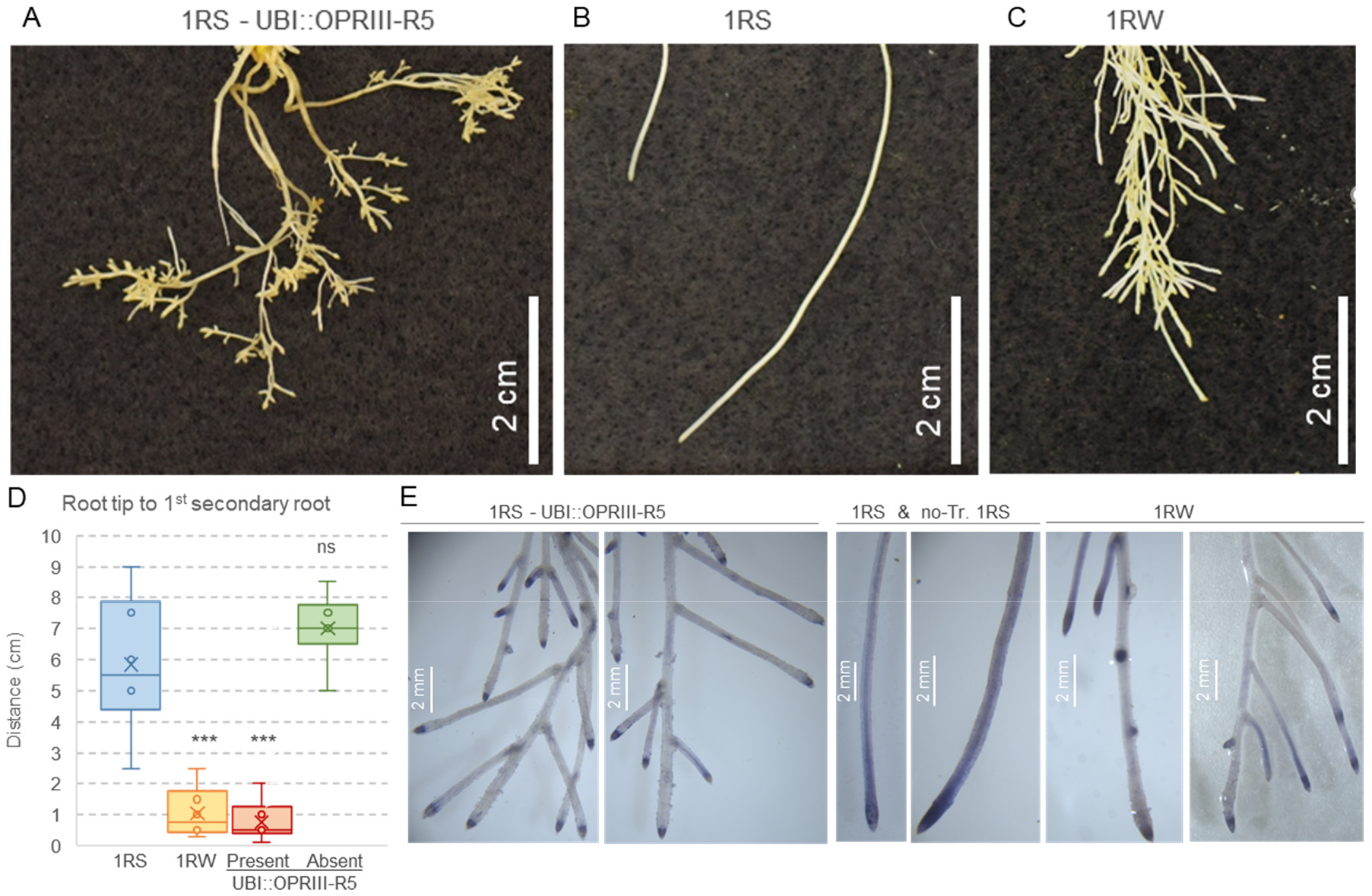
Effect of over-expression of *OPRIII* genes on root architectures. (**A-C**) Last 5 cm of the seminal roots showing different branching patterns. (**A**) UBI::OPRIII-R5, (**B**) 1RS and (**C**) 1RW. (**D**) Distance between the root tip and the first secondary root in 1RS, 1RS^RW^ and UBI::OPRIII-R5. ns = not significant, and *** = *P*< 0.001 based on Dunnett tests relative to 1RS. Raw data and statistics are available in Data S12. (**E**) Roots of the same genotypes as in A at the last time point stained with NBT to visualize the distribution of ROS. 1RS = WT Hahn, no-Tr. 1RS = sister line without the transgene.

We confirmed that wheat *OPRIII-B1*, *OPRIII-B2*, *OPRIII-B3*, and *OPRIII-A2* encode functional 12-oxophytodienoate reductase enzymes. We expressed the proteins and showed that all four genes can reduce 12-oxo-phytodienoic acid (OPDA), 13-epi-12-OPDA and 4,5-ddh-JA substrates using NADPH (Fig. 4A-C, Data S13) ^17,25^. We also measured JA in the terminal 1 cm of the seminal roots at 6 DAG and observed a 2.5 to 3.3-fold increase in JA concentration in 1RW and a 4.8-fold increase in UBI::OPRIII-R5 relative to 1RS (Fig. 4D-E, Data S14). These results demonstrate that increases in *OPRIII* gene dosage or expression are associated with increases in JA in early seminal root development.

**Fig. 4.**
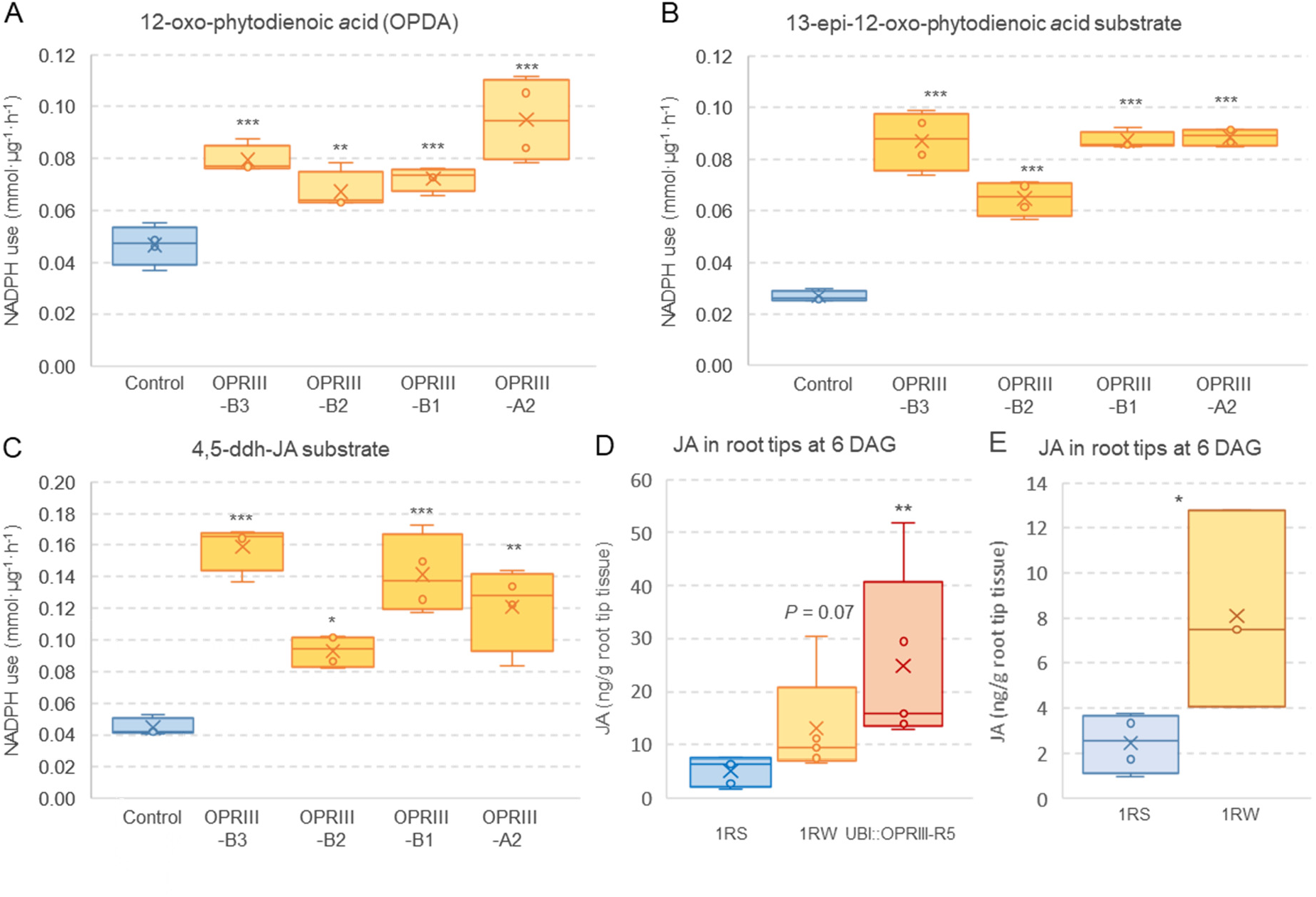
OPRIII role in the biosynthesis of JA. (**A-C**) Consumption of NADPH (nicotinamide adenine dinucleotide phosphate) by different OPRIII proteins using different substrates. (**A**) 12-oxo-phytodienoic acid (OPDA), (**B**) 13-epi-2-oxo-phytodienoic acid (**C**) 4,5-ddh-JA. A boiled mixture of recombinant proteins was used as the negative control. (**D**) Jasmonic acid (JA content) in root tips of 1RS, 1RW and UBI::OPRIII-R5 at 6 DAG. (**E**) Independent experiment for JA content in root tips of 1RS and 1RW at 6 DAG. ns = not significant, * = *P*< 0.05, ** = *P*< 0.01, and *** = *P*< 0.001 based on *t*-tests in E and in Dunnett tests relative to 1RS or control in A-D. n= 4 in all experiments except D where n=5. Raw data and statistics are available in Data S13-14.

To test if the differences in JA-Ile were responsible for the reduced seminal root growth of the 1RW lines, we used Ibuprofen (IBU), a known inhibitor of the JA-Ile biosynthetic pathway ^26^. The addition of IBU-5 μM in the hydroponic culture at six DAG resulted in similar seminal root elongation in 1RS and 1RW (Fig. 5A, Data S15). The addition of IBU-5 μM also accelerated seminal root elongation in 1RW when added at 8 or 10 DAG, but not at 12 DAG (Fig. 5B, Data S16), suggesting that IBU is no longer effective when the JA-Ile signaling cascade is already induced. The addition of IBU-5 μM also eliminated the differences in the distribution of ROS between 1RW and 1RS (Fig. 5C). These results indicate that differences in both root length and ROS distribution between 1RS and 1RW were likely mediated by changes in JA. These two effects may be interconnected since a previous study in Arabidopsis has shown that changes in ROS distribution play an important role in root stem cell maintenance ^27^.

**Fig. 5.**
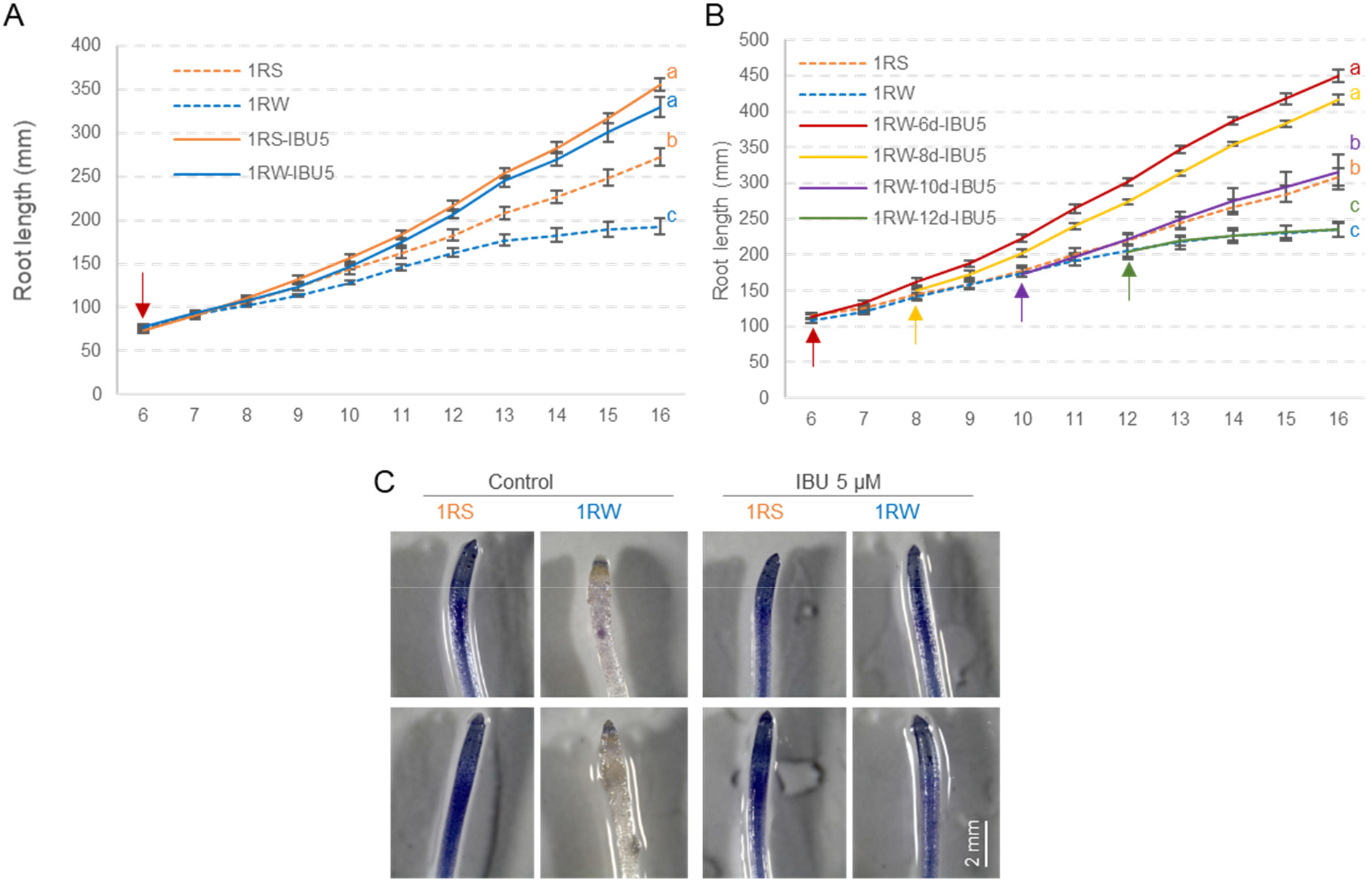
Effect of Ibuprofen (IBU) on root growth. (**A**) Average root length of 1RS and 1RW (n =9-10) with and without IBU 5 μM added at day 6 (red arrow). (**B**) Effect of IBU 5 μM added at 6, 8, 10, and 12 DAG (red arrows) on 1RW root length (n = 8). (**A** and **B**) Different letters indicate significant differences at 16 DAG based on Tukey tests. Error bars are s.e.m. Raw data and statistics are in Data S15-16. (**C**) Effect of IBU 5 μM on ROS distribution along roots (determined with NBT).

A comparison of the 1RS root transcriptomes with those of 1RW and UBI::OPRIII-R5 (Data S3-5, Extended Data Fig. S6) revealed a large numbers of differentially expressed genes (DEGs) at both 6 DAG (5,352 genes) and 16 DAG (4,701 genes, Fig. 6A-B, Data S17-18). These results indicate major developmental changes in the distal region of the seminal roots. Three lines of evidence indicate similar changes in the 1RW and UBI::OPRIII-R5 (transformed into 1RS) transcriptomes relative to 1RS (WT). First, we observed a large number of shared DEGs at 6 (613) and 16 DAG (732) (Fig. 6A-B, Data S19-20). Second, we detected a high and significant regression (*P*< 0.001, *R=* 0.76-0.78) between the log_2_ expression ratios including all genes with more than two-fold difference in expression (Fig. 6C-D, Data S21). Finally, we observed significantly higher correlations (*P*<0.0001) between the DEGs of UBI::OPRIII-R5 and 1RW (*R =* 0.9434 at 6 DAG and *R =* 0.9698 at 16 DAG) than between UBI::OPRIII-R5 and 1RS at the same time points (*R =* 0.9275 at 6 DAG and *R*= 0.9587 at 16 DAG) (Extended Data Fig. S7). Taken together, these results indicate that the transcriptome changes in 1RW and UBI::OPRIII-R5 relative to 1RS are similar and likely driven by the increased *OPRIII* expression in these two lines.

**Fig. 6.**
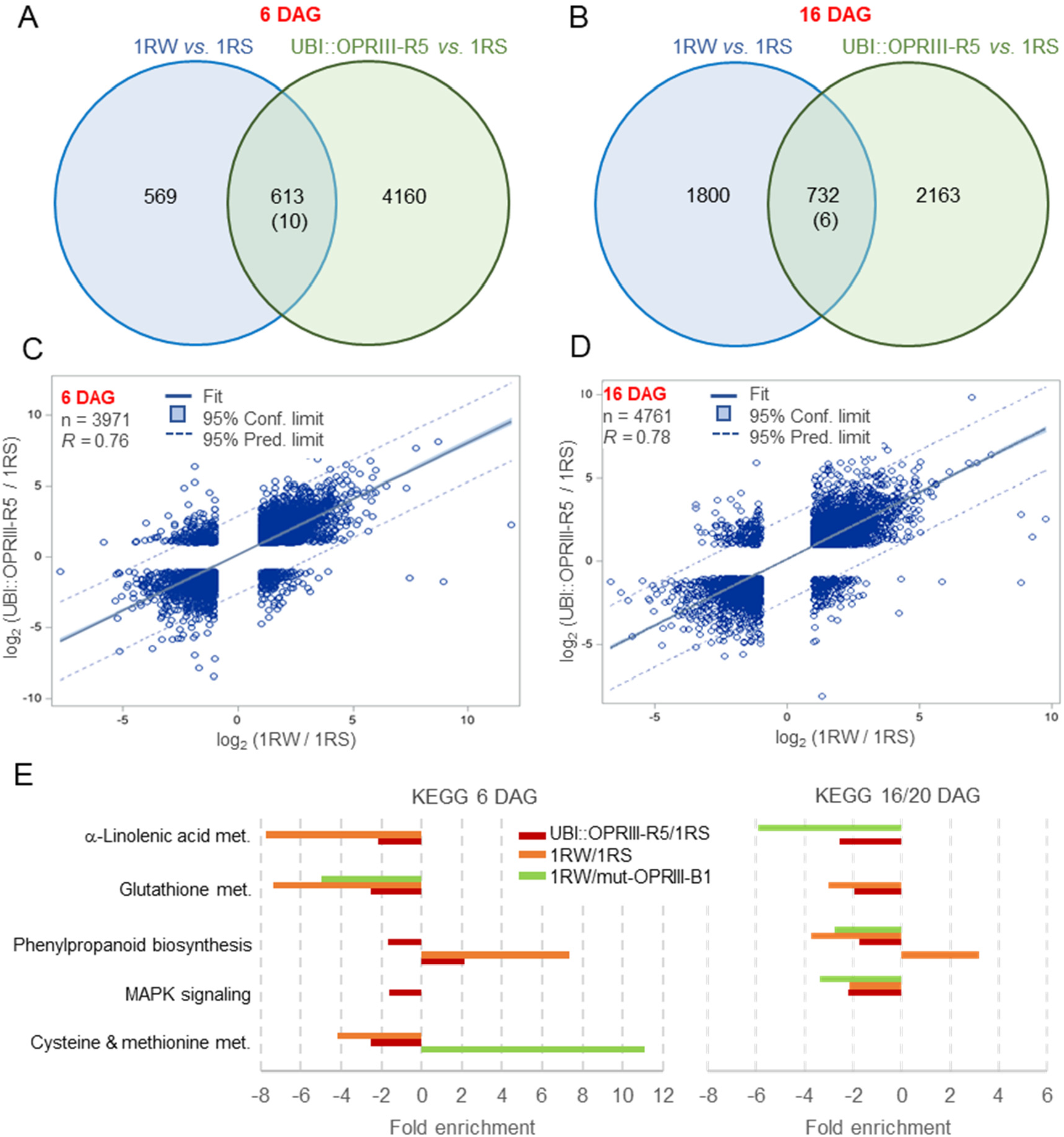
Transcriptome profiles of root tips of Hahn-1RS, 1RW, UBI::OPRIII-R5 and 1RW T_2_ sister lines with and without a 32 bp deletion in *OPRIII-B1*. Samples were collected at 6 for all genotypes and 16 (Hahn-1RS, 1RW, UBI::OPRIII-R5) or 20 days (1RW T_2_ sister lines) after germination (DAG). (**A** and **B**) Differentially expressed genes (DEGs, FDR< 0.05) in comparisons between 1RW *vs.* 1RS and UBI::OPRIII-R5 *vs*. 1RS. The number in the intersection indicates common DEGs in the same direction, and the number in parenthesis in opposite directions (Data S19-20). (**C** and **D**) Regression between log_2_ expression ratios 1RW/1RS and UBI::OPRIII-R5/1RS for genes with more than 2-fold changes in expression relative to 1RS (log_2_ fold < −1 or > +1) and at least one replication >0. Regressions and *R^2^* were calculated based on 3,971 genes in (**C**) and 4,761 genes in (**D**). The significant regressions (*P*< 0.001, Data S21) indicate similar changes in expression in 1RW and UBI::OPRIII-R5 relative to 1RS. **(E)** Significant enriched pathways in KEGG analyses in all three transcriptome comparisons in at least one time point (*P* < 0.05, Data S26). Separate analyses were performed for the early and late time points and for the upregulated (positive values) and downregulated (negative values) DEGs obtained from the comparisons between 1RW and1RS (Data S17), UBI:OPRIII-R5 and 1RS (Data S18), and 1RW T_2_ sister lines with and without a 32 bp deletion in *OPRIII-B1* (Data S24). Transcriptome raw data and statistics are in Data S3-5 and Data S17-26.

To characterize the main pathways affected by the changes in *OPRIII* expression, we carried out an additional Quant-Seq transcriptome analysis comparing RW (WT) and mut-OPRIII-B1(Data S22-24). We performed KEGG analyses (Kyoto Encyclopedia of Genes and Genomes) using the closest rice homologs of the DEGs between the 1RW/1RS and UBI::OPRIII-R5/1RS comparisons at 6 and 16 DAG and for RW/mut-OPRIII-B1 comparisons at 6 and 20 DAG (Data S25-26).These analyses showed three specific pathways that were significantly enriched in all three comparisons (Fig. 6E). The significant enrichment in the alpha-linolenic acid and glutathione pathways are likely associated with the observed changes in the seminal roots in JA (Fig. 4D-E) and ROS (Fig. 5C and previous results ^13^), respectively. An additional pathway significantly enriched at both time points (Fig 6E) was the phenylpropanoid biosynthetic pathway, which is critical for the establishment of root barriers ^28^. The KEGG analyses provide an initial view of the pathways affected in the distal region of the seminal roots by the changes in *OPRIII* expression and JA concentration, but additional functional studies will be necessary to validate the DEGs identified in this transcriptome dataset.

In summary, we identified the *OPRIII* genes as the causal genes for the differences in root length and architecture between the wheat 1RS and 1RW isogenic lines, which were previously associated with significant differences in root depth and grain yield in the field ^12,13^. The extensive variation detected in the number of functional *OPRIII* genes in the available sequenced wheat genomes (Data S2) suggests that natural variation in these genes may have contributed to the adaptation of wheat to different soil environments. The identification of the *OPRIII* gene dosage as a sensitive point in the JA biosynthetic pathway provides a previously unknown target to engineer root architecture in wheat and possibly other cereal crops.

## Acknowledgments

We thank the UC Riverside Metabolomics Core Facility for their help with the Jasmonic Acid determinations and the UC Davis transformation facility for the Fielder transgenic plants. We also thank Yanpeng Wang for his help with the transformation vectors; Rudi Appels for his help with the 1RS gene names in AK58; and Neelima Sinha and Kristina Zumstein for their assistance with the amplicon sequencing to detect induced Crispr-CAS9 mutations.

## Author contributions

G.G. performed most of the experimental work and data analysis, supervised HW, and wrote the first version of the manuscript. JZ, GSM, JM, GFB, and JD contributed data analyses. HW, JZ, TH, contributed to the experimental work. JM and GSM performed the Ibuprofen and jasmonic acid sensitivity experiments. BS, MJC and JT contributed the Hahn transgenic plants. HK and KD performed the JA determinations. GLZ and JYG determined the enzymatic activity of the OPRIII proteins. AL contributed the diiso-1RS and other valuable genetic resources, and MH synthesized and provided the 4,5-ddhJA. JD, TF, and GSM wrote the grant proposals that supported this work. All authors reviewed the manuscript. JD and GSM initiated the project and supervised students and postdocs. JD contributed to the statistical analyses and was responsible for the final manuscript.

## Data and materials availability

Raw data and statistics supporting all figures, supplemental figures and transcriptome analyses are provided in Data S1-26. RNA-seq data was deposited in GenBank under BioProject numbers PRJNA819072 (Hahn-1RS), PRJNA819073 (Hahn-1RW) and PRJNA819075 (Hahn-UBI::OPRIII-R5). Quant-Seq Data was deposited in GenBank under BioProject numbers PRJNA847262 (Hahn-1RW) and PRJNA847590 (mut-OPRIII-B1). Read statistics, expression data for all wheat and rye 1RS genes, and a complete list of differentially expressed genes are provided in the Supplemental online data. The genetic stocks used in this study have been deposited in the National Small Grain Collection as PI 672837, PI 672838, and PI 672839.

## Online content

Supplementary methods, figures and tables are available in the Supplementary Materials

## Supplementary Materials

Online Methods

Extended Data Figures S1 to S7

Supplementary Data S1 to S26

References. Manuscript: 1-28, Extended Data Figures 1-5, and Online Methods 1-22.

## References

1. Zampieri, M., Ceglar, A., Dentener, F. & Toreti, A. Wheat yield loss attributable to heat waves, drought and water excess at the global, national and subnational scales. Environmental Research Letters 12, 064008 (2017).

2. Gupta, A., Rico-Medina, A. & Cano-Delgado, A.I. The physiology of plant responses to drought. Science 368, 266–269 (2020).

3. Pinto, R.S. & Reynolds, M.P. Common genetic basis for canopy temperature depression under heat and drought stress associated with optimized root distribution in bread wheat. Theoretical and Applied Genetics 128, 575–585 (2015).

4. Langridge, P. & Reynolds, M.P. Genomic tools to assist breeding for drought tolerance. Current Opinion in Biotechnology 32, 130–135 (2015).

5. Ehdaie, B., Layne, A.P. & Waines, J.G. Root system plasticity to drought influences grain yield in bread wheat. Euphytica 186, 219–232 (2012).

6. Villareal, R.L., Banuelos, O., Mujeeb-Kazi, A. & Rajaram, S. Agronomic performance of chromosomes 1B and T1BL.1RS near-isolines in the spring bread wheat Seri M82. Euphytica 103, 195–202 (1998).

7. Waines, J.G. & Ehdaie, B. Domestication and crop physiology: roots of green revolution wheat. Annals of Botany 100, 991–998 (2007).

8. Ehdaie, B., Whitkus, R.W. & Waines, J.G. Root biomass, water-use efficiency, and performance of wheat rye translocations of chromosomes 1 and 2 in spring bread wheat 'Pavon'. Crop Science 43, 710–717 (2003).

9. Fenn, D., Lukow, O.M., Bushuk, W. & Depauw, R.M. Milling and baking quality of 1BL/1RS translocation wheats. I. Effects of genotype and environment. Cereal Chemistry 71, 189–195 (1994).

10. Graybosch, R.A. Uneasy unions: Quality effects of rye chromatin transfers to wheat. Journal of Cereal Science 33, 3–16 (2001).

11. Lukaszewski, A.J. Manipulation of the 1RS.1BL translocation in wheat by induced homoeologous recombination. Crop Science 40, 216–225 (2000).

12. Howell, T. et al. Mapping a region within the 1RS.1BL translocation in common wheat affecting grain yield and canopy water status. Theoretical and Applied Genetics 127, 2695–2709 (2014).

13. Howell, T. et al. A wheat/rye polymorphism affects seminal root length and yield across different irrigation regimes. Journal of Experimental Botany 70, 4027–4037 (2019).

14. Li, G.W. et al. A high-quality genome assembly highlights rye genomic characteristics and agronomically important genes. Nature Genetics 53, 574–584 (2021).

15. Rabanus-Wallace, M.T. et al. Chromosome-scale genome assembly provides insights into rye biology, evolution, and agronomic potential. bioRxiv, 2019.12.11.869693 (2019).

16. Gabay, G. et al. Structural rearrangements in wheat (1BS)-rye (1RS) recombinant chromosomes affect gene dosage and root length. Plant Genome 14, e20079 (2021).

17. Li, W.Y. et al. Comparative characterization, expression pattern and function analysis of the 12-oxo-phytodienoic acid reductase gene family in rice. Plant Cell Reports 30, 981–995 (2011).

18. Chen, Q. et al. The basic helix-loop-helix transcription factor MYC2 directly represses *PLETHORA e*xpression during jasmonate-mediated modulation of the root stem cell niche in Arabidopsis. Plant Cell 23, 3335–3352 (2011).

19. Kim, J.M. et al. Acetate-mediated novel survival strategy against drought in plants. Nature Plants 3, 17097 (2017).

20. Mielke, S. et al. Jasmonate biosynthesis arising from altered cell walls is prompted by turgor-driven mechanical compression. Science Advances 7(2021).

21. Wang, S.C. et al. Lateral root formation in rice (*Oryza sativa*): promotion effect of jasmonic acid. Journal of Plant Physiology 159, 827–832 (2002).

22. Li, W.Y. et al. Phylogenetic analysis, structural evolution and functional divergence of the 12-oxo-phytodienoate acid reductase gene family in plants. BMC Evolutionary Biology 9, 90 (2009).

23. Mou, Y.F. et al. Genome-wide identification and characterization of the *OPR* gene family in wheat (*Triticum aestivum* L.). International Journal of Molecular Sciences 20(2019).

24. International Wheat Genome Sequencing Consortium. Shifting the limits in wheat research and breeding using a fully annotated reference genome. Science 361, eaar7191 (2018).

25. Chini, A. et al. An OPR3-independent pathway uses 4,5-didehydrojasmonate for jasmonate synthesis. Nature Chemical Biology 14, 171–178 (2018).

26. Staswick, P.E., Huang, J.F. & Rhee, Y. Nitrogen and methyl jasmonate induction of soybean vegetative storage protein genes. Plant Physiology 96, 130–136 (1991).

27. Yamada, M., Han, X.W. & Benfey, P.N. RGF1 controls root meristem size through ROS signalling. Nature 577, 85–88 (2020).

28. Andersen, T.G. et al. Tissue-autonomous phenylpropanoid production is essential for establishment of root barriers. Current Biology 31, 965–977 (2021).

