## Supplemental Figures and Methods for "Dosage differences in *12-OXOPHYTODIENOATE REDUCTASE* genes modulate wheat primary root growth"

### Extended Data Figures

**Figure S1.** Phylogenetic relationship among OPRIII proteins in rye, wheat and rice. We inferred the evolutionary history of OPRIII proteins from rye, wheat and rice using the Neighbor-Joining method. We used the MUSCLE protein alignment presented in the next page to calculate the optimal tree with branch lengths proportional to the distances used to infer the phylogenetic tree. The evolutionary distances were computed using the Poisson correction method, and are in the units of the number of amino acid substitutions per site. We show the percentage of replicate trees in which the associated taxa clustered together in the bootstrap test (1000 replicates) next to the branches. We removed all ambiguous positions for each sequence pair (pairwise deletion option) and the final dataset included 401 positions. We conducted all the analyses using MEGA X (*2*). OPR nomenclature is based on Mou et al. 2019 (*1*). These authors also provide a more complete phylogenetic analysis of all wheat OPR proteins. We used this analysis to assign names to the rye proteins based on their relationship with the named wheat genes (in red). Gene accession numbers, protein sequences and expression levels are available in Data S2.

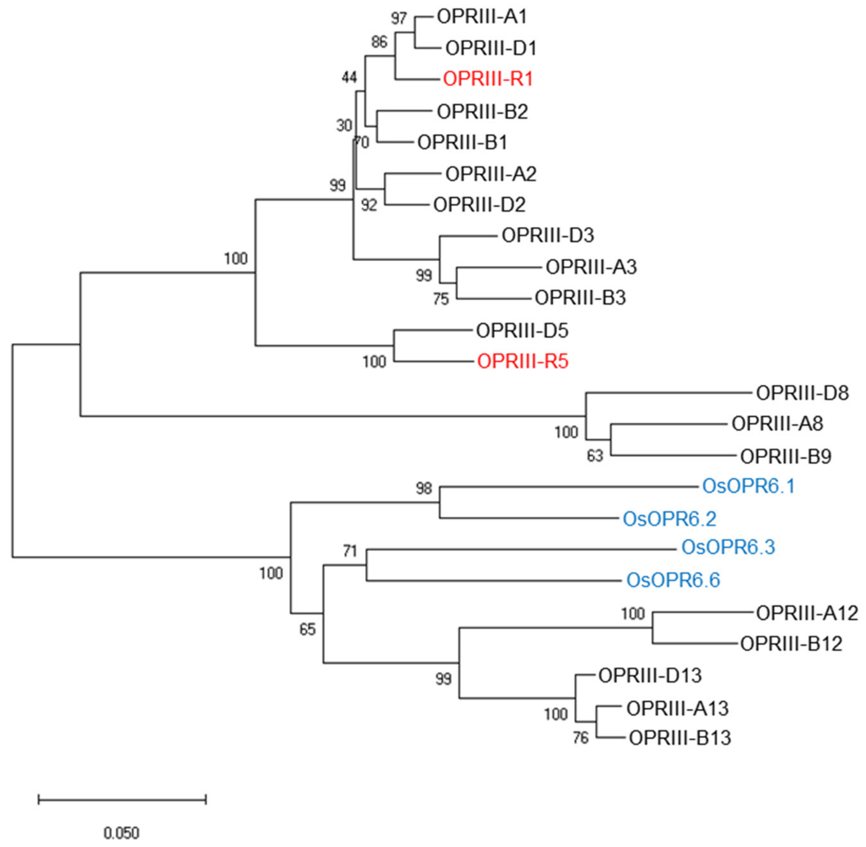


Notes:
**OPRIII-B2** is more closely related with OPRIII1 than with OPRIII2 proteins, and may need to be renamed. OPRIII genes in 1BS are organized in a different order than in 1AS or 1DS suggesting some rearrangements.

**OPRIII-B9** is closely related to the OPRIII8 proteins and may need to be renamed.

OPRIII12 and OPRIII13 located in wheat homeologous group 7 clustered with rice OsOPR6 genes in a colinear region on rice chromosome 6. The other wheat and rye OPRIII genes located in the duplicated region in the short arm of homoeologous group 1 form a separate cluster with no orthologs in rice. A similar clustering has been described in (*1*)

Tree in Newick machine-readable format:

```
((((((( OPRIII-A1,OPRIII-D1),OPRIII-R1),(OPRIII-B2,OPRIII-B1)),(OPRIII-A2,OPRIII-D2)),(OPRIII-D3,(OPRIII-A3,OPRIII-B3))),(OPRIII-D5,OPRIII-R5)),(OPR-D8,(OPR-A8,OPR-B9)),((OsOPR6.1,OsOPR6.2),((OsOPR6.3,OsOPR6.6),((OPRIII-A12,OPRIII-B12),(OPRIII-D13,(OPRIII-A13,OPRIII-B13))))));
```

Alignment of *Triticum aestivum* and *Oryza sativa* (Os) OPRIII proteins using the MUSCLE algorithm in MEGA X. The alignment was visualized using pyBoxshade (https://github.com/mdbaron42/pyBoxshade/blob/master/BS_app.py). Rice protein names are indicated in blue and the closest wheat proteins on homeologous group 7 are indicated in green. Amino acids straddling two exons (encoded by a codon split by an intron) are indicated in red. The wheat OPRIII genes located in homeologous group 7 share the same exon structure and position of straddled amino acids as the rice genes. OPRIII1, OPRIII2 and OPRIII3 proteins have three exons whereas OPRIII5, OPRIII8 and OPRIII9 have two exons. All five OPRIII genes on the short arm of homeologous group 1 share the same border between the last two exons.

```
             ....:...10....:...20....:...30....:...40....:...50....:...60....:...70....:...80....:
OPRIII-A3    ------------MEPIPLLTPYK-----MGQFDLAHRVVLAPLTRRRSYANVPQPHAAVYYSQRATAGGLLIAEATVVSDTGRGY
OPRIII-B3    ------------MEPIPLLTPYK-----MGQFDLAHRVVLAPLTRRRSYANVPQPHAAVYYSQRATAGGLLIAEATVISDTGRGY
OPRIII-D3    ------------MEPIPLLTPYK-----MGQFDLAHRVVLAPLTRRRSYANMPQPHAAVYYSQRATAGGLLIAEATVVSNTGRGY
OPRIII-A2    ------------MEPIPLLTPYK-----MGQFDLAHRVVLAPLTRQRSYGNVPQPHAAVYYSQRATAGGLLIAEATGVSDTAQGY
OPRIII-D2    ------------MEPIPLLTPYK-----MGQFDLAHRVVLAPLTRQRSYGNVPQTHAAVYYSQRATTGGLLIAEATGVSDTAQGY
OPRIII-B2    ------------MEPIPLLTPYK-----MGQFDLAHRVVMAPLTRQRSYGNVPQPHAAVYYGQRATAAGLLIAEATGVSDTAQGY
OPRIII-B1    ------------MEPIPLLTPYK-----MGHFDLAHRVVLAPLTRQRSYGNVPQLHAAVYYSQRATAGGLLIAEATGISDTAQGY
OPRIII-A1    ------------MEPIPLLTPYK-----MGQFDLAHRVVLAPLTRQRSYGNVPQPHAAVYYSQRATAGGLLIAEATGVSDTAQGY
OPRIII-D1    ------------MEPIPLLTPYK-----MGQFDLAHRVVLAPLTRQRSYGNVPQSHAAVYYSQRATAGGLLIAEATGVSDTAQGY
OPRIII-R1    ------------MEPIPLLTPYK-----MGQFDLAHRVVMAPLTRQRSYGNVPQPHAAVYYSQRATAGGLLIAEATGVSDTAQGY
OPRIII-D5    ------------MEPTPLLTPYK-----MGQLDLAHRIVLAPLTRQRSYGNVPQPHAAVYYSQRATVGGLLITEATGVSDTAQGY
OPRIII-R5    ------------MEPIPLLTPYK-----MGQLDLAHRIVLAPLTRQRSYGNVPQPHAAVYYSQRASAGGLLITEATGVSDTAQGY
OPRIII-A8    ---MAGEEGETGAAAIPLLAPYRAG---SGELELAHRVVLAPLTRQRSPGNLPQPHAAVYYAQRATAGGMLITEATGVSAAAQGH
OPRIII-D8    MAGEDGETVTVAAAAIPLLTPYRTG---GGELELAHRVVLAPLTRQRSPGNLPQPHAAVYYAQRATAGGMLITEATGVSAAAQGH
OPRIII-B9    ------------------------------------------MTRQRSPGNLPQPHAAVYYAQRATAGGMLITEATGVSAAAQGH
OPRIII-A12   ---------MATKEPIPLLTPHK-----MGQFELSHRVVLAPLTRCRSYANVPQPHAALHYSQRATKGGLLIAEATGVSATAQGF
OPRIII-B12   ---------MATKEPIPLLTPHK-----MGQFELSHRVVLAPLTRCRSYGNVPQPHAALHYSQRATKGGLLIAEATGVSATAQGF
OPRIII-A13   ---------MVAKEAIPLLTPHR-----MGRFELSHRVVLAPLTRCRSYANVPQPHAAAYYSQRATKGGLLIAEATGVSATAQGY
OPRIII-B13   ---------MVAKEAIPLLTPHR-----MGRFELSHRVVLAPLTRCRSYANVPQPHAALYYSQRATEGGLLIAEATGVSATAQGY
OPRIII-D13   ---------MVAKEAIPLLTPHK-----MGQFELSHRVVLAPLTRCRSYANVPQPHAAAYYSQRATRGGLLIAEATGVSATAQGY
OsOPR6.1     MVQHHQAAANDDHQAIPLLTPYKQAGRPGSKLDLSHRVLLAPMTRCRSYGNVPQPHAALYYTQRATRGGLLITEATGVSATAQGY
OsOPR6.2     ---------MVNQAAMPLLTPYKQA---GGKIDLSHRVLLSPMTRCRSYGNVPQPHAALYYTQRATSGGLLITEATGVSDTAQGY
OsOPR6.3     -MAREAEKDAAAAAEIPLLTPYK-----MGRFELSHRVVLAPLTRNRSYGNVPRPHAVLYYTQRATSGGLLVTEATGVSDTAQGY
OsOPR6.6     --MVHAPAKVAAAAAIPLLTPYK-----MGQLELSHRVVLAPLTRCRSYGNVPQPHAAVYYSQRATRGGLLIAEATDISPTAQGY

             ...90....:..100....:..110....:..120....:..130....:..140....:..150....:..160....:..170
OPRIII-A3    TDTPGIWTAEHVEAWKPIVTAVHAKGALFFCQLWHVGRVSTFELQPGGAA-----PLSSTEKGVGPQLSFDGRLEEFSPPRRLTV
OPRIII-B3    TDTPGIWTAEHVEAWKPIIAAVHAKGALFFCQLWHVGRVSTFELQPGGAA-----PLSSTEKGVGPQMTFDGRLEEFSPPRRLTV
OPRIII-D3    TDTPGIWTAEHVEAWKPIVAAVHAKGALFFCQIWHVGRVSTFELQPGGAA-----PLSSTEKGVGPQMSFDGRLEEFSPPRRLTV
OPRIII-A2    TDTPGIWTTEHIEAWKPIVAAVHAKGALFFCQIWHVGRVSTFELQPGGAA-----PLSSTEKGVGPQISFDGHREEFSPPRRLTV
OPRIII-D2    TDTPGIWTAEHVEAWKPIVAAVHAKGALFFCQIWHVGRVSTFELQPGGAA-----PLSSTEKGVGPQMSFDGHREEFSPPRRLTV
OPRIII-B2    TDTPGIWTAEHVEAWKPIVAAVHAKGALFFCQIWHVGRVSTFELQPGGAA-----PLSSTEKGVGPQMSFDGRLEEFSPPRRLTV
OPRIII-B1    TDTPGIWTAEHVEAWKPIVAAVHAKGALFFCQIWHVGRVSTFELQPGGAA-----PLSSTEKGVGPQMSFDGRLEEFSPPRRLTV
OPRIII-A1    NDTPGIWTAEHVEAWKPIVAAVHQKGALFFCQIWHCGRVSTFELQPGGTA-----PLSSTEKGVGPQMSFDGRLEEFAPPRRLTV
OPRIII-D1    NDTPGIWTAEHVEAWKPIVAAVHQKGALFFCQIWHCGRVSTFELQPGGTA-----PLSSTEKGVGPQMSFDGRLEEFAPPRRLTV
OPRIII-R1    NDTPGIWTAEHVEAWKPIVAAVHAKGALFFCQIWHCGRVSTFELQPGGTA-----PLSSTEKGVGPQMSFDGRLEEFAPPRRLTV
OPRIII-D5    RDTPGVWTAEHVEAWKPIVDAVHAKGALIFCQIWHVGRVSTYEYQPGGAA-----PLSCTDKGVGPQMSYDGRLEEFAPPRRLKV
OPRIII-R5    RDTPGVWTAEHVEAWKPIIDAVHAKGAIIFCQIWHVGRVSTYEYQPGGAA-----PLSCTDKGVGPQMSYDGRLEEFAPPRRLTV
OPRIII-A8    RPTPGVWTDEQVDAWRPVVDAVHAKGAVIFCQLWHVGRVVGK-LRPDGTPAGTPRPVSSTGRPIATPRMNDGVEEEFATPRRLDV
OPRIII-D8    RPTPGVWAGEQVEAWRPVVDAVHAKGAVVFCQLWHVGRVVGK-LRPDGARAETPQPVSSTGRPIATPRMNDGVEEEFATPRRLDV
OPRIII-B9    RPTPGVWTDEQVEAWRPIVDAVHTKGAVIFCQLWHVGRVVGA-LRPDGTRAETV-PVSSTGRPIATPRMNDGVEEEFATPRRLAV
OPRIII-A12   PDTPGIWTQQQVNAWKPIVDAVHSKGALFFCQIWHAGRVSSNEFQPDGHA-----PISSTDKQIPPDAE-SGMV--YSKPRRLHT
OPRIII-B12   PDTPGIWTQQQVDAWKPIVDAVHGKGALFFCQIWHAGRVSTNEFQPDGHA-----PISSTDKQIPPDAE-SDTV--YSKPRQLHT
OPRIII-A13   PETPGIWTQQQVDAWKPIVDAVHRKGALFFCQIWHVGRVSTNDFQPDGQA-----PISSTDKQITPDAE-SGMV--YSKPRQLQT
OPRIII-B13   PETPGIWTQQHVDAWKPIVDAVHRKGALFFCQIWHVGRVSTNDFQPDGQA-----PISSTDQQITPDAE-SGMV--YSKPRQLQT
OPRIII-D13   PETPGIWTQQQVDAWKPIVDAVHRKGALFFCQIWHVGRVSTNDFQPDGQA-----PISSTDKQITPDAE-SGMV--YSKPRQLQT
OsOPR6.1     PETPGVRTREHVEAWKPIVDAVHRKGALFICQLWHVGRVSNNGFQPDGLA-----PISSTDKAITPDGY--GMV--YSKPRRLRT
OsOPR6.2     PETPGVWTREHVEAWKPIVDAVHRKGALFICQLWHVGRVSTNDYQPNGQA-----PISSSDIQITPDGS--GIV--YSKPRRLRV
OsOPR6.3     PDTPGIWTQQQVEAWKPIVDAVHRKGALFICQLWHVGRVSTNEYQPDGQA-----PISSTDRQITPDDS--GIV--YSKPRRLRT
OsOPR6.6     PETPGIYTQQQIEAWKPIVDAVHRKGALFFLQIWHVGRVSTTDFQPNGQA-----PISSTDKQITPDDS--GMV--YSKPRRLRT
```

```
            ....:..180....:..190....:..200....:..210....:..220....:..230....:..240....:..250....:
OPRIII-A3   EEIPAIVDDFRKAARNAIDAGFDGVEIHAANGYIIEQFLKDSANDRTDEYGGSLENRCRFALEVVDAVVKEVGGHRVGIRLSPFT
OPRIII-B3   QEIPAIVDDFRKAARNAIDAGFDGVEIHAANGYIIEQFLKDSANDRTDEYGGSLENRCQFALEVVDAVVKEAGGHRVGIRLSPFA
OPRIII-D3   EEIPALVDDFRKAARNAIDAGFDGVEIHGANGYIIEQFLKDSANDRTDEYGGSLENRCRFALEVVDAVVKEVGGHRVGIRLSPFA
OPRIII-A2   EEIPAIVDDFRKAARNAIDAGFDGVEIHGANGYIIEQFLKDSANDRADEYGGTLENRCRFALEVVDAVVKEVGGRRVGIRLSPFT
OPRIII-D2   EEIPAIVDDFRKAARNAIDAGFDGVEIHGANGYIIEQFLKDSANDRTDEYGGSLENRCRFALEVVDAVVKEVGGRRVGIRLSPFT
OPRIII-B2   EEIPAIIDDFRKAARNAIDAGFDGVEIHGANGYIIEQFLKDSANDRTDEYGGSLENRCRFALEVVDAVVKEVGGHRVGIRLSPFT
OPRIII-B1   EEIPAIIDDFRKAARNAIDAGFDGVEIHGANGYIIEQFLKDSANDRTDEYGGSLENRCRFALEVVDAVVKEVGGHRVGIRLSPFT
OPRIII-A1   EEIPAIVDDFRKAARNAIDAGFDGVEIHGANGYIIEQFLKDSANDRIDEYGGNLENRCRFALEVVDAVVKEVGGHRVGIRLSPFT
OPRIII-D1   EEIPAIVDDFRKAARNAIDAGFDGVEIHGANGYIIEQFLKDSANDRIDEYGGSLENRCRFALEVVDAVVKEVGGHRVGIRLSPFT
OPRIII-R1   EEIPAIVDDFRKAARNVIDAGFDGVEIHGANGYIIEQFLKDSANDRTDEYGGSLENRCRFALEVVDAVIKEVGGHRVGIRLSPFT
OPRIII-D5   EEIPAIVDDFRKAARNAINAGFDGVEIHGANGYLIEQFLKDSANDRTDKYGGSIENRSRFALEVVDAVVKEVGGHRVGIRLSPFT
OPRIII-R5   EEIPAIVDDFRKAARNAIDAGFDGVEIHGANGYLIEQFLKDSANDRNDEYGGSIENRCRFALEVVDAVVKEIGGHRVGIRLSPFT
OPRIII-A8   AEIAGVVDDFRRAARNAVDAGFDGVEIHGAHGYLVEQFLKDSANDRDDEYGGSLENRCRFALEVVTAVAREIGGRRVGVRLSPFA
OPRIII-D8   AGIAGVVDDFRRAARNAVDAGFDGVEIHGAHGYLVEQFLKDSANDRDDEYGGSLENRCRFALEVVTAVAEEIGGRRLGVRLSPFA
OPRIII-B9   EEIAGVLDDFRRAARNAVYAGFDGVEIHGAHGYLVEQFLKDSTNDRDEEYGGSLENRCRFVLEVVTAVADEIGGRRVGVRLSPFA
OPRIII-A12  DEVPLIVDDFRRAARNVIEAGFDGVEIHGAHGFLLEQFMKDSSNDRTDRYGGSLENRCRFAVEVIDAIVHDIGADRVGIRLSPFT
OPRIII-B12  DEVPLIVDDFRRAARNVIEAGFDGVEIHGAHGFLLEQFMKDSSNDRTDMYGGSLENQCRFAVEVIDAIVHEIGADRVGIRLSPFV
OPRIII-A13  DEIPLIVDDFRRAARNAVEAGFDGVEIHGAHGYLLEQFMKDSSNDRTDEYGGSLENRCRFAVEVIDAIVHEIGADRVGIRLSPFV
OPRIII-B13  DEIPLIVDDFRRAARNAIEAGFDGVEIHGAHGYLLEQFMKDSSNDRTDEYGGSLKNRCRFAVEVIDAIVHEIGADRVGIRLSPFV
OPRIII-D13  DEIPLIVDDFRRAARNAIEAGFDGVEIHGAHGYLLEQFMKDSSNNRTDEYGGSLENRCRFAVEVIDAIVHEIGADRVGIRLSPFV
OsOPR6.1    DEIPQIVDDFRLAARNAVEAGFDGVEIHGANGYLLEQFMKDSSNDRTDEYGGSLENRCRFAVEVIDAVVGEIGAHSVGIRLSPFL
OsOPR6.2    DEIPQIVDDFRLAARNAIEAGFDGVEIHGANGYLLEQFMKDSSNDRTDEYGGSLENRCRFAVEVIDAVVGEIGAHRVGIRLSPFL
OsOPR6.3    EEIPQIIDDFRRAARNAIEAGFDGVEIHGAHGYLLEQFMKDSANDRSDEYGGSLENRCRFVVEVIDAIVAEVGAHRVGIRLSPFI
OsOPR6.6    DEIPQIIDDFRRAARNAIEAGFDGVEIHGAHGYLLEQFMKDSANDRTDEYGGSLENRCRFAVEVIDAVVAEVGAHRVGIRLSPFV

            ..260....:..270....:..280....:..290....:..300....:..310....:..320....:..330....:..340
OPRIII-A3   DYMDCHDSDPHSLAFYMSTKLNDH--DILYIHMIEPRMTIEDGRRVVPKRFLPYREAFKGTFIANGGYDRGEGDKVIVEGYTDLV
OPRIII-B3   DYMDCHDSDPHSLALYMSTKLNDH--DILYIHMIEPRMAIVDGRRVVPKRLLPYRKAFKGTFIANGGYDREEGGKVVVEGYTDLV
OPRIII-D3   DYMDCHDSDPHSLALYMSTKLNDH--DILYIHMIEPRMAIVDGRRVVPKRLLPYREAFKGTFVANGGYDREEGGKVVAEGYTDLV
OPRIII-A2   DYMDCHDSDPHSLALYMSTKLNDH--DILYIHMIEPRMAIVDGRRVVPKRLLPYREAFKGTFIANGGYDREEGGKVVTEGYTDLV
OPRIII-D2   DYMDCHDSDPHSLALYMSTKLNDH--DILYIHMIEPRMAIVDGRRVVPKRLLPYREAFKGTFIANGGYDREEGSKVVTEGYTDLV
OPRIII-B2   DYMDCHDSDPHSLALYMSTKLNGH--GILYIHMIEPRMAIVDGRRVVPKRLLPYREAFKGTFIANGGYDREEGGKVITEGYTDLV
OPRIII-B1   DYMDCHDSDPHSLALYMSTKLNDH--GILYIHMIEPRMAIVDGRRVVPKRLLPYREAFKGSFIANGGYDREEGGKVVTEGYTDLV
OPRIII-A1   DYMECHDSDPHSLALYMSTKLNDH--GILYIHMIEPRMAIVDGRRVVPKRLLPYREAFKGTFIANGGYDREEGGKVVTEGYTDLV
OPRIII-D1   DYMDCHDSDPHSLALYLSTKLNDH--GILYIHMIEPRMAIVDGRRVVPKRLLPYREVFKGTFIANGGYDREEGGKVVTEGYTDLV
OPRIII-R1   DYMDCHDSDPHSLALHMSTKLNDH--GILYIHMIEPRMAIVDGRRVVPKRLLPYREAFKGTFIANGGYDREEGGKVVTEGYTDLV
OPRIII-D5   DYMDCHDSDPHALALHMSTKLNDY--NILYLHMVEPRMAIVDGRRVVPKRLLPYREAFKGTFMANGGYDREEGGKVVAEGYTDLV
OPRIII-R5   DYMDCHDSDPHGLALHMSTKLNDY--NIVYLHMVEPRMAIVDGRRVVPKRLLPYREAFKGTFMANGGYDREEGGKVVGEGYTDLV
OPRIII-A8   DYMDCHDSDPHALALHMATKLNGIPGGILYLHMVEPRMARADGRRVVPKRLRPYREAFGGTFIAAGGYDREEGNKVVGEGYADLV
OPRIII-D8   DYMDCHDSDPHALALHMATKLKGIPEGILYLHMMEPRMARADGRRVVPKRLRPYRDAFGGAFIAAGGYDREEGNKVVREGYTDLV
OPRIII-B9   DYMDCHDSDPHALALLMATKLNNVPGGILYLHMVEPRMARANGRRVVPKRLRPYREAFGGTFIAAGGYDR-EGNKVVGEGYADLV
OPRIII-A12  DYMDCFDSNPHVLGTYMVQQLNKHK-EFLYCHMVEPHTTIVDGRMQISHGLLPFRKAFNGTFIAAGGYDREEGNKVVIDSYTDLV
OPRIII-B12  DYMDCFDSNPHVLGTYMVQQLNKHK-EFLYCHMVEPHMTIVDGHMQIPHRLLPFRKAFNGTFIAAGGYDREEGNKMVIDGYTDLV
OPRIII-A13  DYMDCFDSDPHALGMYMVQQLNKHQ-GFVYCHMVEPRMAIVDGRRQIPHGLLPFRKAFNGTFIAAGGYDREEGNKVVADGYADLV
OPRIII-B13  DYMDCFDSDPHALGMYMVQQLNKHQ-GFVYCHMVEPRMAIVDGRRQIPHGLLPFRKAFNGTFIAAGGYDREEGNKVVADGYADLV
OPRIII-D13  DYMDCFDSDPHALGMYMVQQLNKHQ-GFVYCHMVEPRMAIVDGRRQIPHGLLPFRKAFKGTFIAAGGYDREEGNKVVADGYADLV
OsOPR6.1    DYMDCVDSDPEALGSYMVEQLNKHE-DFLYCHMVEPRMAIVDGRRQIQHGLLPFRKQFNGTFIAAGGYDREEGNKAVADGYADLV
OsOPR6.2    DFMDCVDSDPEALGSYMVEQLNKHE-GFLYCHMVEPRMAIVDGRRQIQHGLLPFRKAFKGTFIAAGGYDREEGNKVIENGYTDLV
OsOPR6.3    DYMDCVDSDPVALGSYMVQQLNKHP-GFLYCHMVEPRMAIVEGRRKITHGLLPFRKLFNGTFIAAGGYDREEGNKVIADGYADLV
OsOPR6.6    DFMDCFDSDPVALGSYMVQQLNKHP-GFLYCHMVEPRMAIIEGRRKIAHGLLPFRKQFNGTFIAAGGYDREEGNKVVADGYADLV

            ....:..350....:..360....:..370....:..380....:..390....:..400.
OPRIII-A3   AFGRLFLANPDLPKRFEVGADLNKYDRMTFYTPDPVIGYTDYPFLE---------------
OPRIII-B3   AFGRLFLANPDLPKRFEVGADLNKYDRMTFYTPDPVIGYTDYPFLE---------------
OPRIII-D3   AFGRLFLANPDLPNRFEVGADLNKYDRMTFYTPDPVIGYTDYPFLE---------------
OPRIII-A2   AFGRLFLANPDLPKRFEVGAELNKYDRTTFYTSDPVVGYTDYPFLE---------------
OPRIII-D2   AFGRLFLSNPDLPKRFEIGAELNKYDRMTFYTSDPVVGYTDYPFLE---------------
OPRIII-B2   AFGRLFLANPDLPKRFEVGAELNKYDRMTFYTSDPIVGYTDYPFLE---------------
OPRIII-B1   AFGRLFLANPDLPKRFEVGAELNKYDRMTFYTSDPVVGYTDYPFLE---------------
OPRIII-A1   AFGRLFLANPDLPKRFEVGAELNKYDRMTFYTSDPVVGYTDYPFLE---------------
OPRIII-D1   AFGRLFLANPDLPKRFEVGAELNKYDRMTFYTSDPVVGYTDYPFLE---------------
OPRIII-R1   AFGRLFLANPDLPKRFEVDAELNKYDRMTFYTSDPVVGYTDYPFLE---------------
OPRIII-D5   SFGRSFLANPDLPKRFEIGAELNKYDRMTFYISDPVVGYADYPFLE---------------
OPRIII-R5   SFGRSFLANPDLPKRFEIGAELNKYDRMTFYISNPVIGYTDYPFLE---------------
OPRIII-A8   AFGRLFLANPDLPKRFEFDAKLNGYNRATFYTSDPVVGYTDYPFLG---------------
OPRIII-D8   AFGRLFLANPDLPKRFELDAELNGYDRATFYTSDPVVGYTDYPFLG---------------
OPRIII-B9   AFGRLFLANPGLPKRFELDAELNKYNRATFYTSDPVVGYTDYPFLG---------------
OPRIII-A12  AYGRLFLANPDLPKRFKLDVPLNKYDRSTFYTQDPVIGYTDYPFLDD--------SNTE--
OPRIII-B12  AYGRLFLANPDLPKRFELDVPLNKYDRSTFYTQDPVIGYTDYPFLND--------SNAE--
OPRIII-A13  AYGRIFLANPDLPKRFELDSPLNKYDRNTFYTQDPVIGYTDYPFLEG-------GSNAE--
OPRIII-B13  AYGRIFLANPDLPKRFELDSPLNKYDRNTFYTQDPVIGYTDYPFLEG--------SNAE--
OPRIII-D13  AYGRIFLANPDLPKRFELDSPLNKYDRNTFYTQDPVIGYTDYPFLEG--------SNAE--
OsOPR6.1    AYGRLFLANPDLPKRFELDAPMNNYDRNTFYTQDPVVGYTDYPFLDEHHHDDDDDSNAPSA
OsOPR6.2    SFGRLFLANPDLPKRFELDAPLNKYDRNTFYTQDPIVGYTDYPFLDE-----DQNNSVADA
OsOPR6.3    AYGRHFLANPDLPKRFAINAPLNKYNRSTFYIQDPVVGYTDYPFLDE----KDEGAATYA-
OsOPR6.6    AYGRLFLANPDLPRRFELDAPLNRYDRSTFYTQDPVVGYTDYPFLEE----IDEESRTTYA
```

**Figure S2. Expression of *OPRIII* genes during root development**

Transcript levels in tips of seminal roots of Hahn-1RS and Hahn-1RW collected from 3 to 15 DAG every 3 days. Rye genes (**A**) *OPRIII-R5* and (**B**) *OPRIII-R1*. Wheat genes (**C**) *OPRIII-A1* + *OPRIII-D1*, (**D**) *OPRIII-A2* + *OPRIII-D2*, and (**E**) *OPRIII-B1* (not present in 1RS). Bars indicate average transcript levels relative to *ACTIN* using the delta Ct method. Error bars are s.e.m. based on four replications at each time point for each genotype. Statistical comparisons between genotypes were perform at each time point using *t*-tests except for *OPRIII-B1*, where all 1RS values are 0 and a non-parametric Kruskal-Wallis test was used. ns = not significant, ** = $P < 0.01$ and *** = $P < 0.001$. Raw data, statistics and primers are available in Data S6-7.

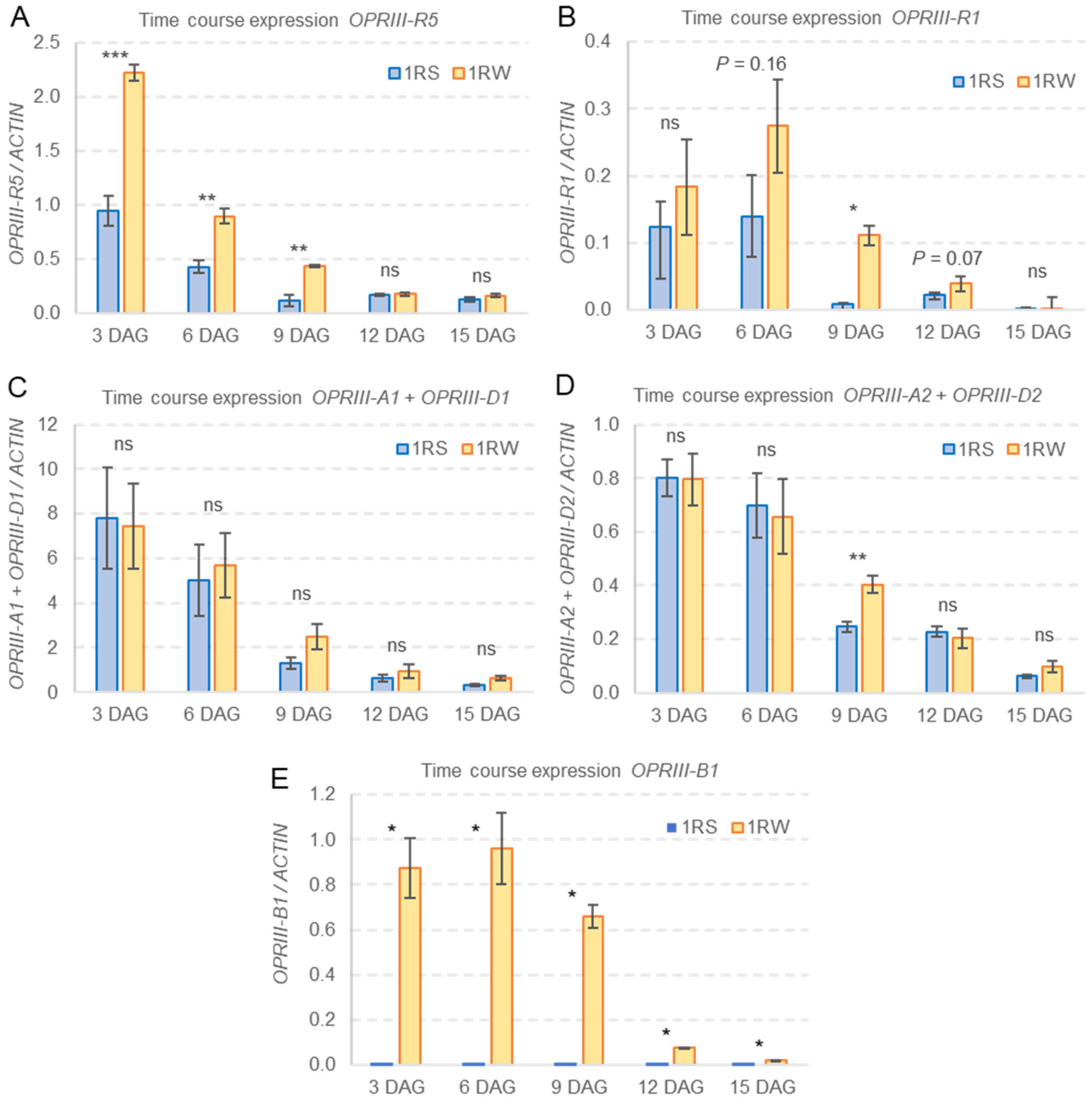

**Figure S3.** Constructs used to generate transgenic Hahn 1RW plants edited for *OPRIII-B1* and Hahn-1RS plants overexpressing wheat *OPRIII-B1* and rye *OPRIII-R5* driven by the maize *UBIQUITIN1* promoter and its intron (UBI1). (**A**) pAct1IHPT-4 (*3*) contains hygromycin phosphotransferase (*hpt*) gene under control of the rice *ACTIN1* promoter, its intron (*act1I*) and the NOS 3' terminator. (**B**) pOsUbi10Cas9 contains the Cas9 gene from pRGEB (*4*) under control of the rice *UBIQUITIN10* promoter, its intron and the NOS 3' terminator. pRGEB32 was a gift from Yinong Yang (http://n2t.net/addgene:63142). (**C**) Guide RNAs Sg51 and Sg52 driven by the *T. aestivum* U6 snRNA gene (GenBank X63066.1) were cloned into vector pENTR/Zeo-H2B (GenBank GU370782). The wheat *OPRIII-B1* (**D**) and rye *OPRIII-R5* genes (**E**) were cloned in the pJIT163-UBI vector (GenBank accession LY758014.1, (*5*)). The *OPRIII* genes were cloned between the maize *UBIQUITIN1* promoter and the CaMV terminator (2,006 to 6,262 bp) replacing the Cas9 region.

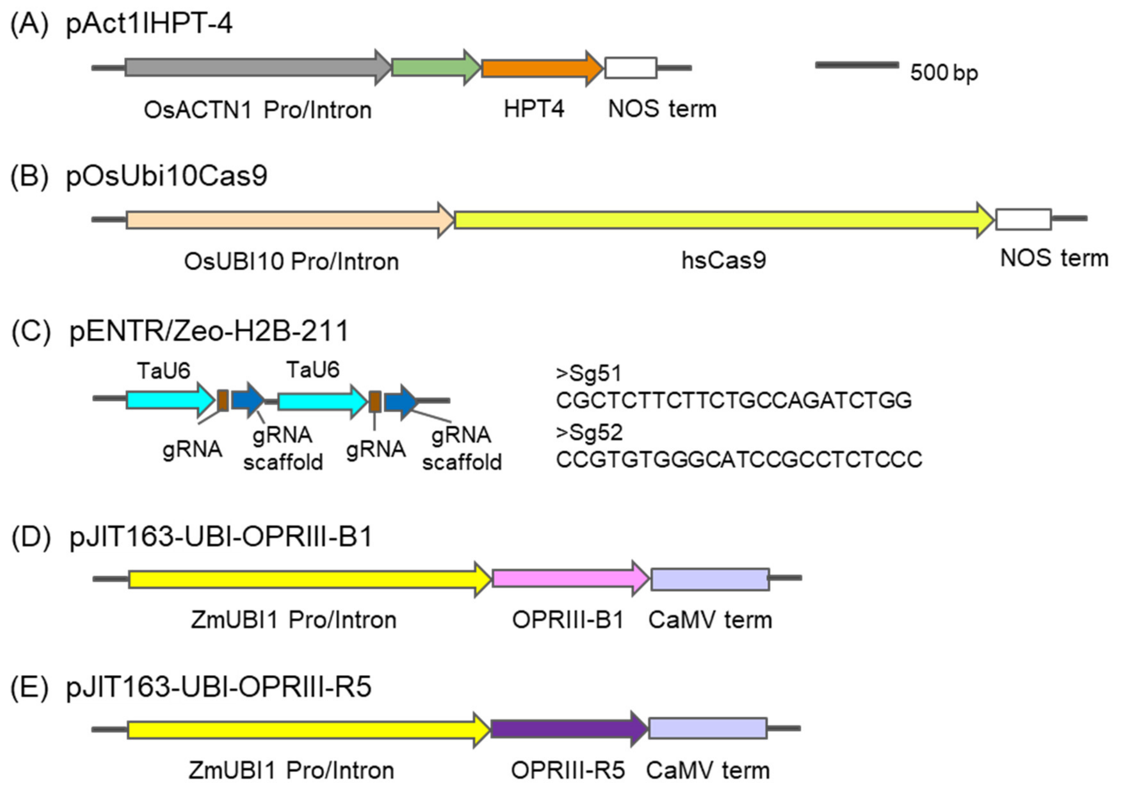

**Figure S4.** Effect of a CRISPR induced 32 bp deletion in *OPRIII-B1* in 1RW $T_3$ sister lines on root length. Time course from 7 to 22 DAG comparing 1RW sister lines with and without the 32 bp deletion in *OPRIII-B1*. The raw data and statistical analyses are presented in Data S8. A similar experiment using $T_2$ sister lines is presented in Fig. 3A in the main text. Error bars are s.e.m. ns = not significant, * = $P< 0.05$ and ** = $P< 0.01$.

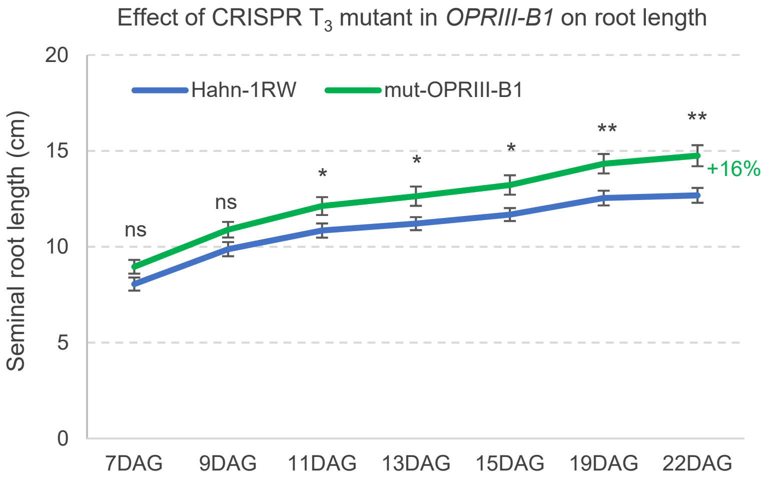

**Figure S5.** Effect of the overexpression of wheat *UBI::OPRIII-B1* and rye *UBI::OPRIII-R5* on root length in wheat cultivar Fielder. Plants carrying the transgene were identified using a combination of an HPT marker in genomic DNA and qRT-PCR markers *UBI::OPRIII-B1* and *UBI::OPRIII-R5* in RNA samples from leaves. (**A**) Time course experiment from 7 to 20 DAG using $T_1$ transgenic plants and sister lines without the transgenes. Asterisks indicate differences between each of the transgenic lines and the wild type using Dunnett tests. ** = $P< 0.01$ and *** = $P< 0.001$. Raw data and statistical tests are available in Data S11. Differences in root length among genotypes were larger and more significant in the later time points, reaching 19% in *UBI::OPRIII-R5* and 25% in *UBI::OPRIII-B1* at 20 DAG. A replicated experiment using $T_3$ plants is presented in the main text in Figure 3D. (**B**) qRT-PCR of *OPRIII-B1* and *OPRIII-R5* relative to *ACTIN* in leaves of $T_3$ *UBI::OPRIII-B1* and *UBI::OPRIII-R5* transgenic plants. Both transgenes are highly overexpressed in the leaves, where the endogenous genes were not detected (Data S11).

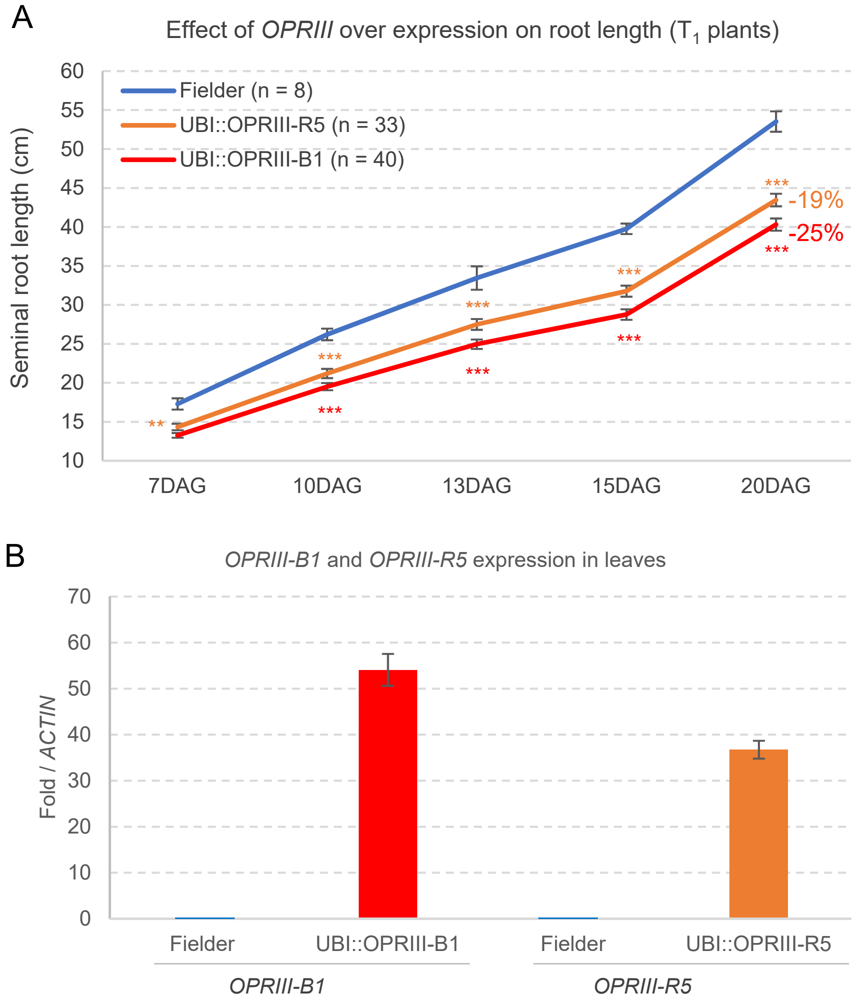

**Figure S6.** Principal component analyses of the 24 RNAseq samples (3 genotypes x 2 time-points x 4 biological replications). (**A**) Analysis based on 8,862 differentially expressed genes (DEGs) between 1RW *vs*. 1RS and between UBI::OPRIII-R5 *vs*. 1RS at both 6 and 16 days after germination (DAG, Data S19). The first principal component (PC1), which explains 61.47% of the variance, mainly separates samples collected at 6 DAG from those collected at 16 DAG (developmentally regulated genes). PC2, which explains 14.41 % of the variation, mainly separates the genotypes within the same day. To visualize better the relationships among genotypes we also performed separate PCA for (**B**) 6 DAG and (**C**) 16 DAG.

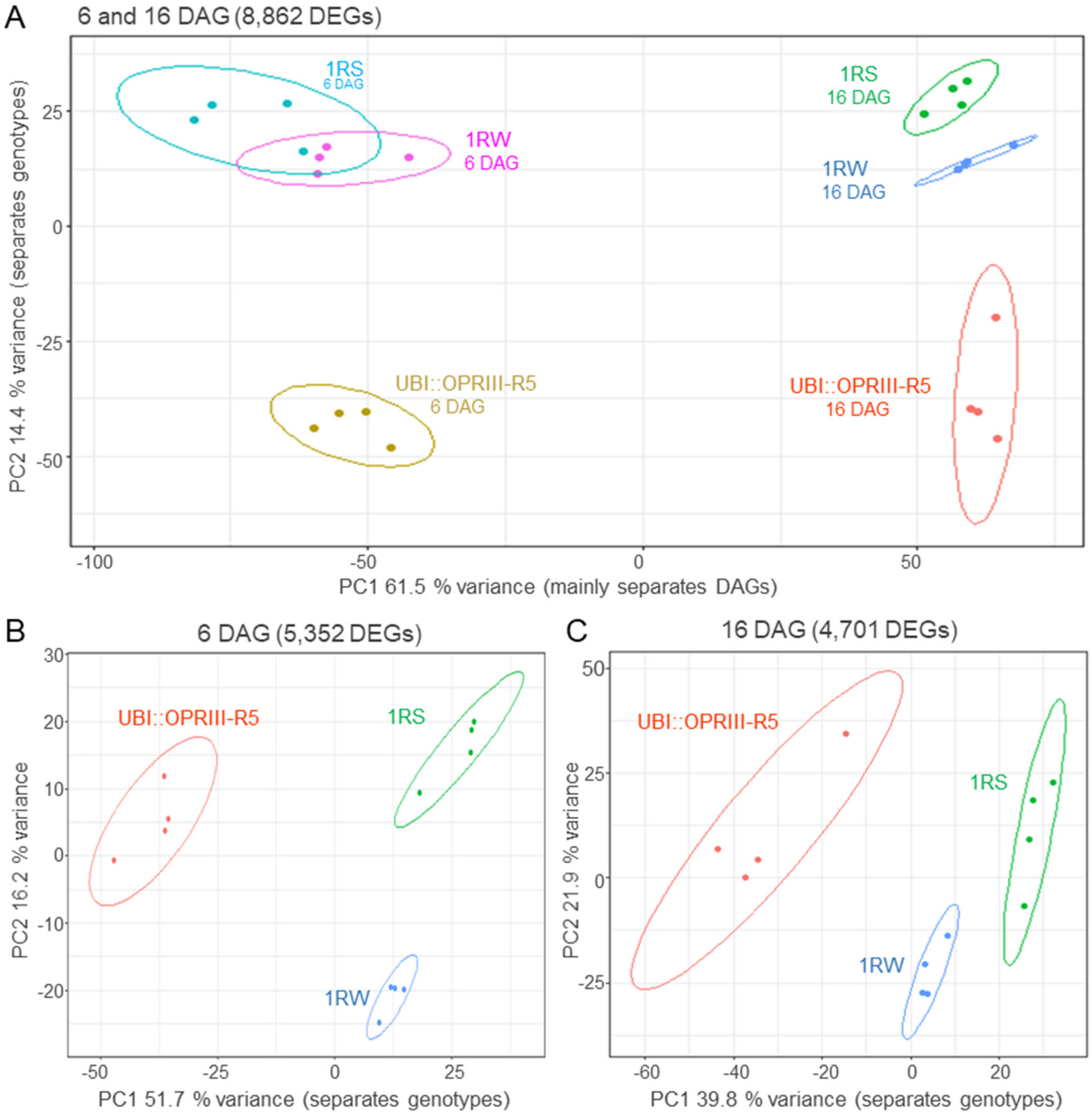

**Figure S7.** Comparisons of transcriptomes of seminal root tips of Hahn-1RS, 1RW, UBI::OPRIII-R5. Regression between TMM values of DEGs genes (n= 8,862, Data S19) genes in 1RW vs. UBI::OPRIII-R5 (A and C), and 1RS vs. UBI::OPRIII-R5 (B and D) at 6 (A and B) and 16 (C and D) days after germination. Significantly higher correlations (*P*<0.0001) between the TMM of the DEGs in UBI::OPRIII-R5 (transformed into 1RS) and 1RW (*R* = 0.9434 at 6 DAG and *R* = 0.9698 at 16 DAG) than between UBI::OPRIII-R5 and 1RS at the same time points (*R* = 0.9275 at 6 DAG and *R*= 0.9587 at 16 DAG). These results indicate more similar changes in expression in 1RW and UBI::OPRIII-R5 relative to 1RS

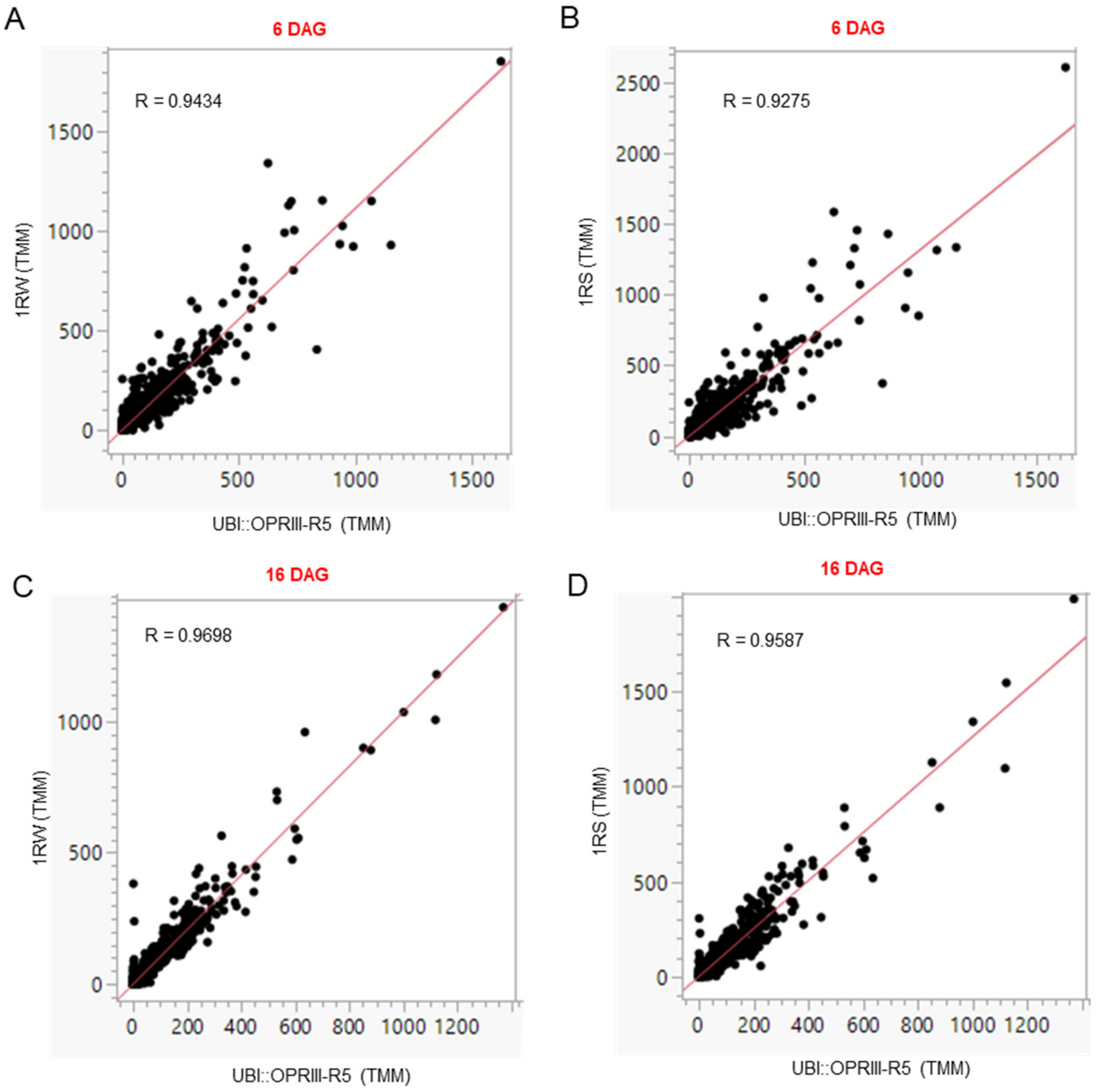

### References


1. Y. F. Mou *et al.*, Genome-wide identification and characterization of the *OPR* gene family in wheat (*Triticum aestivum* L.). *Int. J. Mol. Sci.* **20**, (2019).
2. S. Kumar, G. Stecher, M. Li, C. Knyaz, K. Tamura, MEGA X: Molecular evolutionary genetics analysis across computing platforms. *Mol. Biol. Evol.* **35**, 1547-1549 (2018).
3. M.-J. Cho, W. Jiang, P. G. Lemaux, Transformation of recalcitrant barley cultivars through improvement of regenerability and decreased albinism. *Plant Sci.* **138**, 229-244 (1998).
4. K. Xie, B. Minkenberg, Y. Yang, Boosting CRISPR/Cas9 multiplex editing capability with the endogenous tRNA-processing system. *Proc. Natl. Acad. Sci. U.S.A.* **112**, 3570-3575 (2015).
5. S. Jin *et al.*, Rationally designed APOBEC3B cytosine base editors with improved specificity. *Mol. Cell* **79**, 728-740 (2020).

### Supplemental Methods

#### Hahn-1RS and 1RW isogenic lines

The cultivar Hahn, developed by the "Centro Internacional de Mejoramiento de Maíz y Trigo" (CIMMYT), carries the 1RS.1BL translocation and is referred here as Hahn-1RS. Hahn-1RS was used as recurrent parent for the introgression of two interstitial segments of the 1BS arm from Pavon by homeologous recombination (1WW, PI 672837. Fig. 1A) (*1*). The 1WW chromosome was then backcrossed six times into Hahn-1RS and the isogenicity of the $BC_6F_2$ sister lines was confirmed using the wheat Illumina 90,000 SNP iSelect array (*2*). The Hahn 1WW line was crossed to the original cv. Hahn-1RS to generate a line carrying only the proximal 1BS segment (1WR, PI 672838, for proximal Wheat and distal Rye segments) and another one carrying only the distal 1BS segment (1RW, PI 672839) (Fig. 1A).

#### Generation of diiso 1RS and determination of its effect on root length

An isochromosome 1RS was recovered during manipulation by centric mis-division of the 1RS.1BL translocation in wheat (*3*). This translocation originated from the Kavkaz source of the translocation, via cv. Genaro. After self-pollination, a diisosomic 1RS line of Pavon 76 was isolated. For this study, the diiso 1RS of Pavon 76 was crossed, and backcrossed four times to Hahn-1RS with cytological selection for the presence of diiso 1RS in each generation. The $BC_4F_1$ was self-pollinated and individual plants with diiso 1RS as well as telo 1RS were isolated, grown and self-pollinated. As Hahn-1RS has the 1RS.1BL translocation, the homozygous diiso 1RS addition line (2n= 44) has six doses of the rye chromosome 1RS. A cytogenetic study of the progeny of a diiso 1RS plant showed that this extra diiso chromosome is not stable and is lost in approximately 31% of the progeny (Data S1).

To determine the 1RS copy number variation (CNV) in 23 progenies of a diiso 1RS plant, we performed six independent DNA extractions for each of the 23 plants, and from 8 Hahn-1RS and 8 Hahn-1RW additional plants as controls. We then used qRT-PCR primers qrt1RS5-F and -R (Data S7) to determine the dosage of the *OPRIII-R5* genes located in the 1RS chromosome arm. We used *CO2* as an endogenous control for a single copy gene as reported before (*4*). For each of the 23 progenies and 16 controls, we determined root length in hydroponic tanks as described below, and then calculated a regression between the 1RS CNV and root length using SAS v9.4. To select plants with more than two 1RS chromosomes, we performed *t*-tests between the CNV in each recombinant and the 1RS control, which has two 1RS arms. Root length in plants with more than two 1RS arms were compared with plants carrying the 1RS or 1RW controls. Raw data and statistical analyses for these experiments are available in Data S1.

#### Hydroponic experiments with Ibuprofen

Seeds were imbibed for 3 days at 4 °C and then sowed on a floating mesh in a 0.5 mM $CaCl_2$ solution at room temperature. Four days after, healthy seedlings were transferred to 350 mL pots (one plant per pot) containing an aerated nutrient solution with the following composition:

$Ca(NO_3)_2$ 1.0 mM, KCl 1.5 mM, $KH_2PO_4$ 0.2 mM, $MgSO_4.7H_2O$ 1.0 mM, $CaCl_2$ 1.5 mM, FeEDTA 0.1 mM, $H_3BO_3$ 1 µM, $(NH_4)_6Mo_7O_{24}.4H_2O$ 0.05 µM, $CuSO_4.5H_2O$ 0.5 µM, $ZnSO_4.7H_2O$ 1 µM, $MnSO_4.H_2O$ 1 µM, brought to pH 6.0 ± 0.1 with $Ca(OH)_2$. The solution was renewed three times a week for the duration of the experiment. The experiments were performed in a growth chamber set at 22-23 °C with a photoperiod of 16 h light/8 h dark provided by a fluorescent light source supplemented with incandescent lighting. Photon flux density at the plant level was 150 µmol $m^{-2}$ $s^{-1}$. The length of the second longest seminal root was measured with a ruler four hours after the start of the light period. Ibuprofen sodium salt was added from a SIGMA stock solution (IBU; SIGMA: I1892). Nitroblue tetrazolium (NBT) staining to determine superoxide anion distribution was performed as described before (*5*).

**Stable transformation**

***Transgenic plants generated by particle bombardment*:** Transgenic Hahn-1RS and 1RW plants were transformed via particle bombardment at UC Berkley. Seeds of these two lines were sown weekly and grown in growth chambers. Immature embryos (IEs) were prepared according to the same specifications as Fielder (*6*). They were pre-incubated at 26 °C overnight and used for bombardment. On the day of bombardment, IEs were placed on to DBC3 medium containing mannitol and sorbitol (0.2 M each) (*6, 7*) for osmotic pretreatment. Four hours after treatment with osmoticum, IEs were bombarded as previously described (*7*). Two milligrams of 0.6 µm gold particles were coated with 5 µg of a mixture of pAct1IHPT4 and pJIT163-UBI::OPRIII-B1 or pJIT163-UBI-OPRIII-R5 at a 1: 2 ratio for the transformation of Hahn-1RS (Extended Data Fig. S3). For the transformation of Hahn-1RW, we used the same amount of gold particles coated with 10 µg of a mixture of pAct1IHPT4, pOsUbi10Cas9 and pENTR/Zeo-H2B-211 at a 1:2:3 ratio (Extended Data Fig. S3). pENTR/Zeo-H2B-211 included two guide RNAs targeted to a conserved region in *OPRIII-D3*, *OPRIII-A2*, *OPRIII-A3*, *OPRIII-B1*, and *OPRIII-B2* (sg51*)* and *OPRIII-A2, OPRIII-A1, OPRIII-B1, OPRIII-B2, OPRIII-D5, OPRIII-D2, OPRIII-D1, OPRIII-R1, OPRIII-R5* (sg52, Extended Data Fig. S3). Each particle preparation was resuspended in 85 µL of 100% EtOH and 7.5 µL was spread onto the center of a macrocarrier inside of a macrocarrier holder. The microparticle preps were used for bombardment with a Bio-Rad PDS-1000/He biolistic device (Bio-Rad, Hercules, CA) at 650 psi. Each plate of IEs was bombarded twice per treatment. Sixteen hours post bombardment, IEs were transferred to DBC3 medium and incubated at 26 °C for one week in dim light. Following the resting period, IEs went through 3 rounds of selection via DBC3 media containing 30 mg/L hygromycin B, each round of selection lasting 3 weeks. After the third round of selection, regeneration was initiated using DBC6 media (*8*) containing 30 mg/L hygromycin B and incubated at 26 °C in high light (90 µmol $m^{-2}$ $s^{-1}$) and subcultured every 3 weeks. Once shoots were approximately 0.5-3.0 cm in height, they were transferred to WR rooting media containing 30 mg/L hygromycin B. Plantlets were then transferred to soil once they had enough roots to support transplant to soil.

Hahn-1RW was co-bombarded with vectors pOsUbi10Cas9 (Extended Data Fig. S3) and pENTR/Zeo-H2B-211 including guide RNAs Sg51 and Sg52 (Extended Data Fig. S3). Mutations in the Hahn-1RW plants transformed with the CRISPR-Cas9 vector were screened for mutations in multiple *OPRIII* genes by next-generation sequencing (NGS, MiSeq, Illumina at the UC Davis Genomic Center). Amplicons were obtained as described before (*9*) using nonspecific primers to detect mutations in genes with high identity levels (primers in Data S7). Variants were called for each sample (demultiplexed fastq files) using CRISgo v5 (https://github.com/pinbo/CRISgo). Demultiplexed reads were also mapped to potential targets using BWA v0.7.17 (bwa mem command) (*10*). Variants called from CRISgo were validated by visual inspection of the bam files using IGV v2.7.2 (https://software.broadinstitute.org/software/igv/) (*11*)

***Transgenic plants generated by Agrobacterium-mediated transformation*****:** The hexaploid wheat Fielder transgenic plants were generated at the UC Davis Plant Transformation Facility (http://ucdptf.ucdavis.edu/) using the Japan Tobacco (JT) vector pLC41 (hygromycin resistance) and *Agrobacterium*-mediated transformation technology licensed to UC Davis. The wheat *OPRIII-B1* and rye *OPRIII-R5* were cloned in the binary vector pLC41 (Japan Tobacco) downstream of the maize *UBIQUITIN* promoter and upstream of the 3xHA tag and a NOS terminator. Hygromycin was used as a selectable marker. Transformation was performed using protocols described previously (*12*). Plants were grown in hydroponic tanks as described before (*5*), and root length measurements were performed every two to three days starting at 7 days after germination (DAG). After the last measurement, leaf samples were collected to extract RNA and DNA to identify the transgenic plants using primers described in Data S7.

### qRT-PCR of *OPRIII* genes

The expression levels of several *OPRIII* genes were characterized using quantitative reverse transcription PCR (qRT-PCR) using primers described in Data S7. Hahn-1RS and 1RW plants were grown in hydroponic tanks and the terminal 1 cm of the seminal roots of each plant were collected at different time points (3, 6 ,9, 12, and 15 DAG). Roots from 12 plants were pooled per replication to obtain sufficient RNA, and four pools were used as replications for each time point / genotype combination.

RNA samples were extracted using the Spectrum Plant Total RNA Kit (Sigma-Aldrich). First-strand cDNAs were synthesized from 1 µg of total RNA using the High Capacity Reverse Transcription kit (Applied Biosystems). Quantitative PCR was performed using SYBR Green and a 7500 Fast Real-Time PCR system (Applied Biosystems). *ACTIN* was used as an endogenous control. Transcript levels for all genes are expressed as linearized fold-*ACTIN* levels calculated by the formula $2^{(ACTIN\ \mathrm{CT}\ -\ TARGET\ \mathrm{CT})}$ ± standard error (SE) of the mean, which indicates the ratio between the initial number of molecules of the target gene and the number of molecules of *ACTIN*.

### Determination of OPRIII enzymatic activity

To overcome the redundancy of *OPRIII* homologs and paralogs, we first amplified *OPRIII-B3* and *OPRIII-B1* from 1RW with primers on the UTR region (OPRIII-B3-F / R and OPRIII-B1-F /

R, Data S7). We then cloned the PCR products into T vector and sequenced them to confirm the presence of the complete coding region. We next added the attB site to the coding region using primer OPRIII-B-attB-F combined with either OPRIII-B3-attB-R or OPRIII-B2-attB-R (Data S7). We were not able to clone *OPRIII-B2* and *OPRIII-A2* from the reverse transcription products, so we cloned each exon from genomic DNA using the primers described in Data S7. Using the genomic clones as templates we obtained the full-length coding region by overlap PCR. The *OPRIII* genes were cloned into vector pDONR207 using the Gateway BP Clonase® enzyme and then transformed into pHIS9, a Gateway compatible destination vector modified from pET28a (*13*). The destination plasmids were transformed into *E. Coli*, Rosetta (DE3) for protein expression.

One positive colony per clone was picked into a 10 mL LB liquid medium and cultured overnight at 37 °C. One mL was transferred into 1 liter of LB liquid medium and cultured for ~6 h until $OD_{600}$ 0.6. A stock of isopropyl-β-D-thiogalactoside (IPTG) was added at 1 mM into the cell culture, which was cultivated for 20 h at 16 °C. The cells were collected by centrifugation and stored overnight at -80 °C. The cells were resuspended in 30 mL of lysis buffer (50 mM Tris pH 7.4, 500 mM NaCl 20 mM Imidazole 1% Triton-X100, 1 mM PMSF). The cells were sonicated for 45 min with 5 s sonicate/25 s stop cycles until the sample was clear. The samples were centrifuged at 5,500 g for 10 min at 4°C. The supernatant was centrifuged at 20,000 g for one hour at 4 °C. The supernatant was transferred to a clean 50 mL tube and mixed with 100 µL of Ni-NTA beads Qiagen, Item No. 30210） for 2 h at 4 °C with gentle rotation at 30 rpm. The samples were centrifuged at 500 g for 30 min at 4 °C to collect the pellet. The pellet was washed with 1 mL wash buffer (50 mM Tris pH 7.4, 500 mM NaCl 20 mM Imidazole 1% glycerin) and centrifuged at 500 g for 30 s to remove the supernatant. We repeated the above step twice to remove supernatant contaminations. Pellets were resuspended with 300 µL of elution buffer (50 mM Tris pH 7.4, 500 mM NaCl 250 mM Imidazole 1% glycerin) and kept at 4 °C for 30 min to elute the recombinant protein.

The enzyme activity assay system contained 50 mM PBS, 1 mM NADPH, and 3 µg of the substrate at a 200 µL volume in UV compatible clear 96-well plates (Corning). One µg of recombinant protein was added into the above system to initiate the reaction. OD340 was recorded every minute for 1 h. A boiled mixture of recombinant proteins was used as negative control. A NADPH dilution from 0.25, 0.5, 0.75, 1, and 1.5 mM was used to construct the standard curve. All the samples were analyzed with four replicates.

The 12-oxo-10,15(Z)-phytodienoic acid substrate (CAS No. 85551-10-6) was purchased from Larodan (Item No. 13-1821), whereas the 13-*epi*-12-oxo-phytodienoic acid (CAS No. 71606-07-0) was purchased from Cayman Chemical (Item No. 10195). The synthesis of the 4,5-didehydro jasmonic acid substrate (complete name (+)-3(R),7(S)-4,5-didehydrojasmonic acid, CAS No. 123357-39-1) was reported before (*14*). NADPH was purchased from Real-Times, Beijing Biotechnology (Item No. 041939).

**Jasmonic acid determination**

***Sample preparation*: **Hahn-1RS, 1RW and UBI::OPRIII-R5 plants were grown in hydroponic tanks as described above. At 6 DAG, we collected the terminal one cm of the seminal roots. Eight plants were pooled from each genotype grown in the same tank to collect 100 mg of fresh tissue. Five tanks were used as replications. Samples were frozen and sent to the UC Riverside Metabolomics Core Facility, where the 100-mg samples were ground and mixed with 500 µL of 6:3:1 methyl tert-butyl ether (MTBE):methanol:water with the addition of deuterated internal standards. Samples were vortexed 90 minutes at 4 °C then centrifuged 30 min at 3000 x g and 4 °C, and 200 µL supernatant was transferred to a new 2 mL autosampler vial. Extracts were then dried under nitrogen gas, resuspended in 200 µL methanol, vortexed for 5 minutes to mix, and transferred to an insert for LC-MS analysis.

***Standard Curve*: **A standard curve was prepared by first adding 40 µL of phytohormones standards mix at 10 µg/mL to 760 µL 6:3:1 MTBE:methanol:water with deuterated internal standards, then performing 2-fold serial dilutions. 200 µL aliquots were dried under nitrogen gas, resuspended in 200 µL methanol, vortexed for 5 minutes to mix, and transferred to inserts for analysis.

***LC-MS phytohormones [T3 column]*: **Phytohormone quantitation was performed on a TQ-XS triple quadrupole mass spectrometer (Waters) coupled to an I-class UPLC system (Waters) (*15*). Separations were carried out on a T3 C18 column (2.1 x 100 mm, 1.8 µM) (Waters). The mobile phases were (A) water and (B) acetonitrile, both with 0.1% formic acid. The flow rate was 300 µL/min, and the column was held at 45 °C. The injection volume was 2 µL. The gradient was as follows: 0 min, 0.1% B; 1 min, 0.1% B; 6 min, 55% B; 7 min, 100% B; 8 min, 100% B; 8.5 min, 0.1% B; 13 min, 0.1% B. The MS was operated in selected reaction monitoring mode. Source and desolvation temperatures were 150 °C and 600 °C, respectively. Desolvation gas was set to 1100 L/h and cone gas to 150 L/h. Collision gas was set to 0.15 mL/min. All gases were nitrogen except the collision gas, which was argon. The capillary voltage was 1 kV in positive ion mode and 2 kV in negative ion mode. A quality control sample, generated by pooling equal aliquots of each sample, was analyzed periodically to monitor system stability and performance. Samples were analyzed in random order.

### RNA seq (1RW, 1RS and UBI::OPRIII-R5)

***RNA extraction***: The terminal 1 cm of the three seminal roots of each plant were collected at 6 and 16 DAG. Roots from 12 plants were pooled per replication to obtain sufficient RNA, and four pools were used as replications for each time point/ genotype combination. RNA samples were extracted using the Spectrum Plant Total RNA Kit (Sigma-Aldrich).

***Library preparation for transcriptome sequencing*: **Messenger RNA was purified from total RNA using poly-T oligo-attached magnetic beads. After fragmentation, the first-strand cDNA was synthesized using random hexamer primers, followed by the second strand cDNA synthesis using dTTP for a non-directional library. The library for transcriptome sequencing was ready after end repair, A-tailing, adapter ligation, size selection, amplification, and purification. The library was checked with Qubit and real-time PCR for quantification and bioanalyzer for size distribution detection. The quantified libraries were pooled and sequenced on Illumina platforms. The clustering of the index-coded samples was performed according to the manufacturer's instructions (Novogene). After cluster generation, the library preparations were sequenced on an Illumina platform and paired-end reads were generated. The number of reds per sample and different quality and mapping statistics are described in Data S3.

Reads were mapped to the Chinese Spring Genome RefSeq v1.0 (*16*) combined with the 1RS arm from cultivar AK58 (*17*), allowing a maximum of 1 SNP. Reads were mapped using the splicing aware STAR aligner from the Lexogen pipeline. Reads mapping to more than one location were distributed equally among the identical targets. Expression values were calculated using the trimmed mean of M-values normalization method (TMM, Data S4) (*18*). The sequence of the 1RS.1BL translocation in Aikang58 (AK58) is available only as a preprint and no final gene names have been published, so we provide a table with the different names and genome coordinates to facilitate future cross-reference (Data S5) (*17*)

Reads were deposited in GenBank short reads archive under BioProject numbers PRJNA819072 (Hahn-1RS, https://dataview.ncbi.nlm.nih.gov/object/PRJNA819072?reviewer=2aqmui18cbgtqa85bmukufaid4); PRJNA819073 (Hahn-1RW, https://dataview.ncbi.nlm.nih.gov/object/PRJNA819073?reviewer=t4eleafcbqml1016207pg4otpg); and

PRJNA819075 (Hahn UBI::OPRIII-R5, https://dataview.ncbi.nlm.nih.gov/object/PRJNA819075?reviewer=en6fenic3s5p8ie3c478b1p176).

#### Quant-Seq – 1RW and mut-OPRIII-B1 in RW

#### RNA extraction

We performed the RNA-extraction as described above (RNA-seq). Samples were collected at 6 and 20 DAG. The second collection was done at 20 days (and not at 16 days as the RNA seq) since the differences in root length were more significant at 20 DAG. The effect of a single gene mutation is expected to be smaller than the duplication of multiple *OPRIII* genes in 1RW relative to 1RS or the constitutive expression in UBI::OPRIII-R5.

Sequences from the 16 samples (2 genotypes × 2 developmental stages × 4 biological replicates) were generated using Hi-seq (100-bp reads not paired) at the UC Davis Genome Center. The number of Quant-Seq reds per sample and different quality and mapping statistics are described in Data S22. We processed the raw reads using DOE JGI BBTools (https://sourceforge.net/projects/bbmap/) program bbduk.sh to remove Illumina adapter contamination and low-quality reads (forcetrimleft = 21 qtrim = r trimq = 10). Reads were mapped and TMM values were calculated as described above (RNA-seq, Data S23).

Reads were deposited in GenBank short reads archive under BioProject numbers PRJNA847262 (Hahn-1RW: https://dataview.ncbi.nlm.nih.gov/object/PRJNA847262?reviewer=frmphvrui384bduln7fjrs43gc) and PRJNA847590 (mut-OPRIII-B1: https://dataview.ncbi.nlm.nih.gov/object/PRJNA847590?reviewer=4hqqbm07tp93a8vmiirm5n4mri)

***Differential expressed genes (DEG)***: All successfully mapped reads were subjected to differential expression analysis using the DESeq2 R package (*19*) in comparisons between 1RW *vs.* 1RS and UBI::OPRIII-R5 *vs.*1RS, both at 6 and 16 DAG and 1RW vs. mut-OPRIII-B1 at 6 and 20 DAG. Only transcripts with CPM >0.5 in at least 2 samples were included for the last step of the analyses. Genes with an adjusted *P*-value based on a false discovery rate (FDR< 0.05) were considered significant DEGs.

***Regression between $log_2$(1RW /1RS) and $log_2$ (UBI::OPRIII-R5 / 1RS)***: To explore the similarity between the changes in the root transcriptomes of UBI::OPRIII-R5 and 1RW relative to 1RS, we performed a regression analysis between the $log_2$ fold changes in these two comparisons. We first eliminated all genes with 0 counts in all four reps in any of the genotypes to avoid ratios with 0 as denominator and logs of 0, and then calculated the average TMM for each gene at 6 and 16 DAG for the three genotypes. We determined the $log_2$ of the ratios between the averages in 1RW / 1RS and UBI::OPRIII-R5 / 1RS for 6 and 16 DAG separately and for each day retained only those genes showing changes in expression larger than two-fold for the two ratios (log2 ratio < -1 or > +1). We then performed a regression analysis of and plotted the results using PROC REG in SAS 9.4.

***Principle component analysis (PCA)***: A PCA of the 24 RNA-seq samples (3 genotypes x 2 time-points x 4 biological replications) was carried out with pcaExplorer using 8,862 differentially expressed genes (*20*).

***Extraction and visualization of overlapping genes***: The extraction and visualization of the overlapping DEGs between the comparisons of 1RS vs 1RW and 1RS vs UBI::RS6.4 at 6 and 16 DAG was performed with Venny (https://bioinfogp.cnb.csic.es/tools/venny/index.html).

***Correlation analysis***

Significant differences in correlation of the DEGs (TMM values) between (1RW/ UBI::OPRIII-R5) and (1RS/ UBI::OPRIII-R5) was determined with "cocor" ( https://cran.r-project.org/web/packages/cocor/index.html and a web application; http://comparingcorrelations.org/)

***KEGG analysis***: The DEGs obtained from the comparisons between 1RW *vs.*1RS, UBI:OPRIII-R5 *vs.* 1RS and 1RW vs. mut-OPRIII-B1 were subjected to separate KEGG analyses at 6 DAG and 16 or 20 DAG. The analyses were carried out using DAVID (*21, 22*). Enriched pathways were considered significant using *P*<0.05.

### Methods-Only References


1. A. J. Lukaszewski, Manipulation of the 1RS.1BL translocation in wheat by induced homoeologous recombination. *Crop Sci.* **40**, 216-225 (2000).
2. T. Howell *et al.*, Mapping a region within the 1RS.1BL translocation in common wheat affecting grain yield and canopy water status. *Theor. Appl. Genet.* **127**, 2695-2709 (2014).

3. A. J. Lukaszewski, Reconstruction in wheat of complete chromosome-1B and chromosome-1R from the 1RS.1BL translocation of Kavkaz origin. *Genome* **36**, 821-824 (1993).

4. A. Diaz, M. Zikhali, A. S. Turner, P. Isaac, D. A. Laurie, Copy number variation affecting the *Photoperiod-B1* and *Vernalization-A1* genes is associated with altered flowering time in wheat (*Triticum aestivum*). *PLoS One* **7**, e33234 (2012).

5. T. Howell *et al.*, A wheat/rye polymorphism affects seminal root length and yield across different irrigation regimes. *J. Exp. Bot.* **70**, 4027-4037 (2019).

6. J. Tanaka, B. Minkenberg, S. Poddar, B. Staskawicz, M.-J. Cho, Improvement of gene delivery and mutation efficiency in the CRISPR-Cas9 wheat (*Triticum aestivum* L.) genomics system via biolistics. . *bioRxiv* (2022).

7. M.-J. Cho, C. D. Ha, P. G. Lemaux, Production of transgenic tall fescue and red fescue plants by particle bombardment of mature seed-derived highly regenerative tissues. *Plant Cell Rep.* **19**, 1084-1089 (2000).

8. M.-J. Cho, J. Banh, M. Yu, J. Kwan, T. J. Jones, Improvement of *Agrobacterium*-mediated transformation frequency in multiple modern elite commercial maize *(Zea mays* L.) inbreds by media modifications. *Plant Cell Tiss. Org.* **121**, 519-529 (2015).

9. N. R. Campbell, S. A. Harmon, S. R. Narum, Genotyping-in-Thousands by sequencing (GT-seq): A cost effective SNP genotyping method based on custom amplicon sequencing. *Mol. Ecol. Resour.* **15**, 855-867 (2015).

10. H. Li, R. Durbin, Fast and accurate short read alignment with Burrows-Wheeler transform. *Bioinformatics* **25**, 1754-1760 (2009).

11. J. T. Robinson *et al.*, Integrative genomics viewer. *Nat. Biotechnol.* **29**, 24-26 (2011).

12. J. M. Debernardi *et al.*, A GRF-GIF chimeric protein improves the regeneration efficiency of transgenic plants. *Nat. Biotechnol.* **38**, 1274-1279 (2020).

13. S. Wang *et al.*, YR36/WKS1-mediated phosphorylation of PsbO, an extrinsic member of Photosystem II, inhibits photosynthesis and confers stripe rust resistance in wheat. *Mol. Plant* **12**, 1639-1650 (2019).

14. A. Chini *et al.*, An OPR3-independent pathway uses 4,5-didehydrojasmonate for jasmonate synthesis. *Nat. Chem. Biol.* **14**, 171-178 (2018).

15. A. M. Sheflin *et al.*, High-throughput quantitative analysis of phytohormones in sorghum leaf and root tissue by ultra-performance liquid chromatography-mass spectrometry. *Anal. Bioanal. Chem.* **411**, 4839-4848 (2019).

16. International Wheat Genome Sequencing Consortium, Shifting the limits in wheat research and breeding using a fully annotated reference genome. *Science* **361**, eaar7191 (2018).

17. Z. Ru *et al.*, 1RS.1BL molecular resolution provides novel contributions to wheat improvement. *bioRxiv*, 2020.2009.2014.295733 (2020).

18. M. D. Robinson, A. Oshlack, A scaling normalization method for differential expression analysis of RNA-seq data. *Genome Biol.* **11**, (2010).

19. M. I. Love, W. Huber, S. Anders, Moderated estimation of fold change and dispersion for RNA-seq data with DESeq2. *Genome Biol.* **15**, (2014).

20. F. Marini, H. Binder, pcaExplorer: an R/Bioconductor package for interacting with RNA-seq principal components. *BMC Bioinformatics* **20**, 331 (2019).

21. D. W. Huang, B. T. Sherman, R. A. Lempicki, Systematic and integrative analysis of large gene lists using DAVID bioinformatics resources. *Nat. Protoc.* **4**, 44-57 (2009).

22. D. W. Huang, B. T. Sherman, R. A. Lempicki, Bioinformatics enrichment tools: paths toward the comprehensive functional analysis of large gene lists. *Nucleic Acids Res.* **37**, 1-13 (2009).
